# mrHARDIflow : A pipeline tailored for the preprocessing and analysis of Multi-Resolution High Angular diffusion MRI and its application to a variability study of the PRIME-DE database

**DOI:** 10.1101/2021.11.22.469616

**Authors:** Alex Valcourt Caron, Amir Shmuel, Ziqi Hao, Maxime Descoteaux

## Abstract

The validation of advanced methods in diffusion MRI requires finer acquisition resolutions, which is hard to acquire with decent Signal-to-Noise Ratio (SNR) in humans. The use of Non-Human Primates (NHP) and anaesthesia is key to unlock valid microstructural maps, but tools must be adapted and configured finely for them to work well. Here, we propose a novel processing pipeline implemented in Nextflow, designed for robustness and scalability, in a modular fashion to allow for maintainability and a high level of customization and parametrization, tailored for the analysis of diffusion data acquired on multiple spatial resolutions. Modules of processes and workflows were implemented upon cutting edge and state-of-the-art MRI processing technologies and diffusion modelling algorithms, namely Diffusion Tensor Imaging (DTI), Constrained Spherical Deconvolution (CSD) and DIstribution of Anisotropic MicrOstructural eNvironments in Diffusion-compartment imaging (DIAMOND), a multi-tensor distribution estimator. Using our pipeline, we provide an in-depth study of the variability of diffusion models and measurements computed on 32 subjects from 3 sites of the PRIME-DE, a database containing anatomical (T1, T2), functional (fMRI) and diffusion (DWI) imaging of Non-Human Primate (NHP). Together, they offer images acquired over a range of different spatial resolutions, using single-shell and multi-shell b-value gradient samplings, on multiple scanner vendors, that present artifacts at different level of importance. We also perform a reproducibility study of DTI, CSD and DIAMOND measurements outputed by the pipeline, using the Aix-Marseilles site, to ensure our implementation has minimal impact on their variability. We observe very high reproducibility from a majority of diffusion measurements, only gamma distribution parameters computed on the DIAMOND model display a less reproducible behaviour. This should be taken into consideration when future applications are performed. We also show that even if promising, the PRIME-DE diffusion data exhibits a great level of variability and its usage should be done with care to prevent instilling uncertainty in statistical analyses.

## 1 INTRODUCTION

Magnetic resonance imaging is a modality of medical imaging that has seen an exponential increase of interest in the past few decades since its the only non-invasive technology able to acquire both functional and structural information from soft tissues. That makes it the preferential choice for the study of brain functions and connectivity with the aim, notably, of mapping the human connectome. However, the study of the generated images is riddled with challenges that the scientific community has yet to overcome [O’Donnell and Pasternak (2015)][Jones et al. (2013)][Jones (2010)], especially when it comes to tractography [Schilling et al. (2021b)][Rheault et al. (2020)][Calamante (2019)][Maier-Hein et al. (2017)][Schilling et al. (2017)], even more in ensemble studies using multiple sites and multiple scanner vendors [Schilling et al. (2021a)][Andica et al. (2020)][De Santis et al. (2019)]. This is particularly true for dMRI, for which the current equipment available still fails to provide the intense and fast changing magnetic fields required for good SNR; quality of the produced images is restricted by eddy currents and phase related distortions, to name a few. Still, all MRI related techniques are limited by resolution orders of magnitude above the size of the tissues of interest, resulting in partial volume effects at the interface between mediums. Lowering the size of acquired voxels requires a longer acquisition time, and is often not a viable possibility, since it leads to increased thermal noises in the reception coils and a higher probability of motion-related artifacts in the resulting images. Selecting a good set of acquisition parameters is thus a multi-objective optimization problem, a trade-off between the image resolution required for subsequent modelling and the acceptable level of noise, the duration of the imaging sequence and the type of subject, a research setting versus a clinical application, amongst others. Up to this day, most of it is done by trial and error, based on recommendation from previous experiments, due to the lack of gold standards defining the validity of most empirical MRI measurements. Inasmuch, the absence of standardized processing pipelines, carefully designed to operate universally or given images acquired with a good subset of parameters, further increases the difficulty of generating good research material exempt of biases from the equipment, subject or even processing train.

This validation problem, affecting the field from the MRI scanner parametrization all the way to modelization and quantity measurement, has been extensively discussed and a plethora of techniques have been devised to tackle it. Comparing MRI acquisitions to tracers studies are known to provide good benchmarks in white matter, but can lack in specificity [Heilingoetter and Jensen (2016)] and its application requires the injection of an invasive substance in the tissue of interest thus being only performed ex-vivo. Quite recently, multi-resolution frameworks have gained in popularity, combining low-resolution dMRI to high-resolution microscopy in order to refine the angular representation of diffusion [Howard et al. (2019)]. However, this technique is also limited to ex-vivo studies, since microscopy requires slicing and staining of the brain tissues.

Another good technique that has made its marks is comparison to results in small animals, since they enable the use of specialized equipment that operates at higher magnetic fields and of stronger techniques to mediate with subject motion and cardiac and respiratory gating, which are either inaccessible or ethically impossible in human studies. Those allow to go to finer resolutions, both spatial and angular, while retaining good SNR. Nonetheless, images still require to be adequately preprocessed before fitting models and producing measurements. The algorithms used to that extent must be finely tuned to their specific requirements. While multiple processing pipelines have been designed by the research community [Cieslak et al. (2021)][Autio et al. (2020)][Theaud et al. (2020)][Gorgolewski et al. (2011)][Pierpaoli et al. (2010)], integrating MRI processing technologies into push-button applications, a great deal of them are configured to handle human subjects images or images acquired using a predefined set of parameters specific to a study, and running them on non-human high-resolution diffusion imaging requires extensive re-configuration. Most of them are also offered as black-boxes, difficult to customize or extend, which makes adapting them to different studies time consuming.

The pipeline we consider in this work alleviates most, if not all of those problems. Implemented in Nextflow [Tommaso et al. (2017)], with the latest implementation of their DSL2 framework, we designed a collection of processing modules consisting of pipeline processes and workflows, each targeting a specific set of preprocessing, model reconstruction, quantity measurement or utility algorithms, using cutting edge DWI processing technologies, such as Dipy [Garyfallidis et al. (2014)], FSL [Smith et al. (2004)], Mrtrix [Tournier et al. (2019)] and ANTs [Avants et al. (2008)]. We then built a complete processing pipeline using those modules, tailored to handle multi spatial resolution and high angular resolution diffusion encoding. The use of Nextfow ensures its scalability on a wide range of computing infrastructures, from an execution on a local computer to multiple nodes in an HCP facility, and container technologies (Docker and Singularity [Kurtzer et al. (2017)]) allow for the encapsulation of all dependencies and their versioning. This combination allow for a robust and automated execution, while keeping installation and maintenance as simple as possible for the end-user.

To test the pipeline, we processed 32 datasets from 3 sites of the PRIME-DE [Neff (2019)][Milham et al. (2018)] database. Those contain diffusion weighted images with multiple spatial and angular resolutions, single-shell and multi-shell b-value samplings, multiple scanner vendors and artifacts at different ranges of intensities depending on the site where the acquisition was executed. We provide a brief reproducibility study of the pipeline results, as well as an in-depth variability study of diffusion modelling and measurements obtained from its execution.

## 2 METHOD

### 2.1 Processing library

3 core concepts were taken into account for the development of the processing library for this project :

1. Efficiency High resolution images are heavy (hundreds of megabytes to a few gigabytes), in particular for diffusion MRI, where a complete MRI volume is acquired for each gradient direction. Combining high spatial and orientational resolution leads to image files weighing multiple gigabytes and a number of voxels of the order of several hundreds of millions. Processing must thus be efficient and parallelized as much as possible. To achieve such a degree of efficiency, we coded the pipeline structure using Nextflow [Tommaso et al. (2017)], a meta-scheduling language enabling automatic optimization and execution of a processing tree over multiple datasets and multiple processing nodes.
2. Modularity and Morphability State-of-the-art methods in diffusion MRI are still changing rapidly. For a good pipeline to stay up to date, it requires being able to morph to new standards and integrate novel techniques being discovered. Using the DSL2 framework of Nextflow [Tommaso et al. (2017)], we produced a highly modular library of processes and workflows, used as building blocks in our pipeline. This allowed for a high level architecture in the final pipeline coded for the current study, with a finer description of the dataflows encapsulated in the modules referring to specific processing steps. Using this paradigm makes it easier for new pipelines to be developed that only change a subset of processing steps or add new workflows on top of older ones.
3. Usability and Reproducibility The libraries used for diffusion MRI are ever-changing. It is thus impossible to guarantee a robust execution of our pipeline on versions of its dependencies released in the future. Docker and Singularity [Kurtzer et al. (2017)] - two technologies enabling the encapsulation of libraries in portable packages - were used to wrap up the numerous applications called in the different processing steps of the pipeline. They enable locking the versions of the dependencies into a pre-packaged image and remove the need for the end-user to install and manage them. In addition, they allow for multi-stage building of images, which makes it easy to add new dependencies and modify a set of versions in the case of an update of the underlying pipeline or for the needs of a specific study.

To address these concepts, the library was fragmented into 4 nested and interlocked scopes : pipeline scope, input scope, process scope and workflow scope, as depicted in figure 1. Of them, only the process scope contains the actual calls to algorithms and their input requirements. It encapsulates the actual workhorse of the pipeline and forms the processes module.

**Figure 1.**
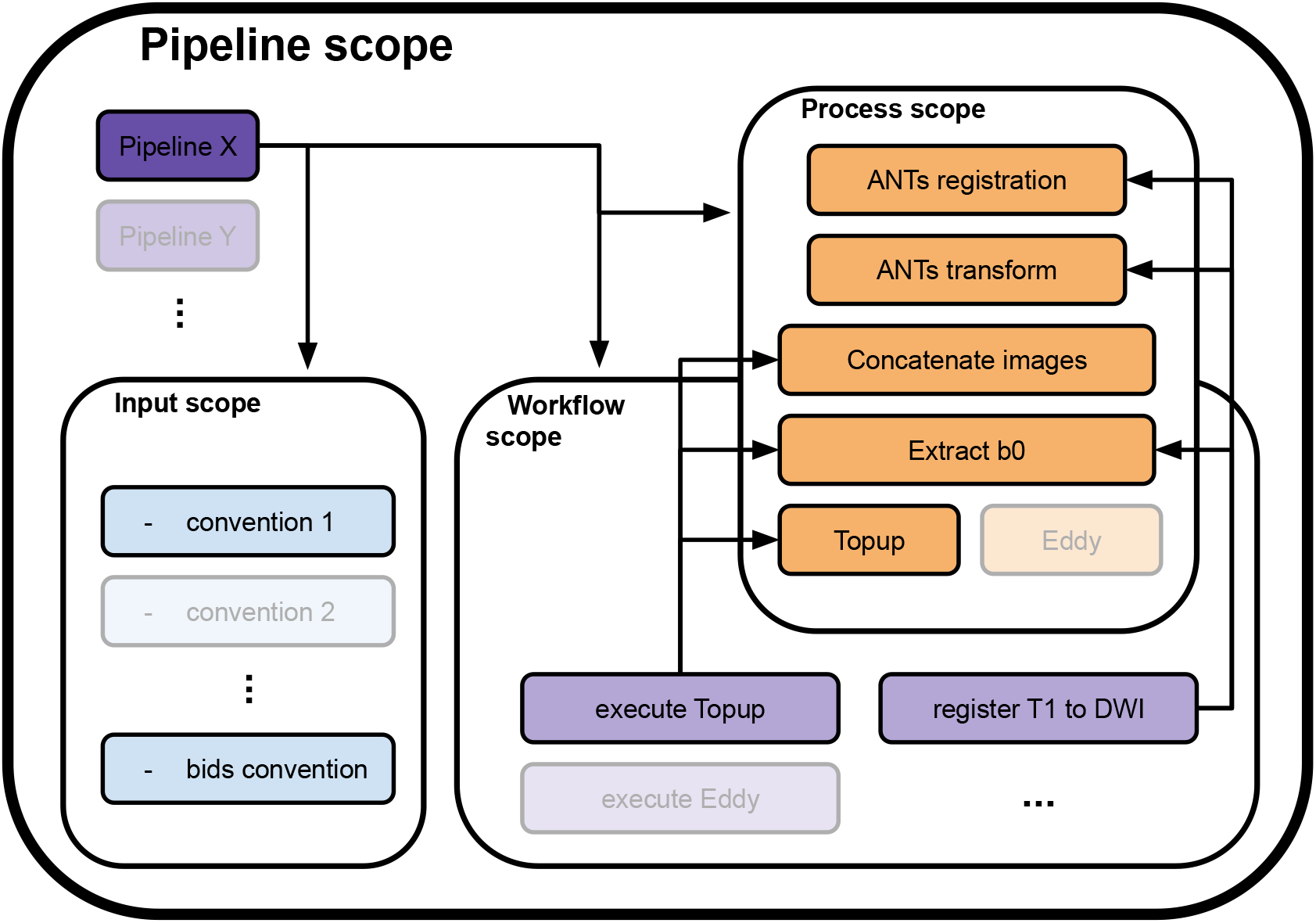
4 scopes composing the Nextflow library and how they interface with each others. A pipeline definition (Pipeline X for example) is placed at the root of the library. To load the data, it uses one or more convention from the input scope (convention 1 and the bids convention). Then, to process it, it calls utilities in either or both the process scope (ANTs registration, Concatenate images and others) and the workflow scope (execute Topup, then register T1 to DWI). A call to an utility in the workflow scope will trigger calls to processes in the process scope they include to process the data (execute Topup firsts calls a concatenation of two opposite phase encoding DWI and extracts their b0 volumes, to pass the result to Topup for phase distortions correction). The input, workflow and process scopes form the core of the Nextflow library. Their content can be used by any pipeline defined in the pipeline scope, allowing high levels of code reusability.

**Figure 2.**
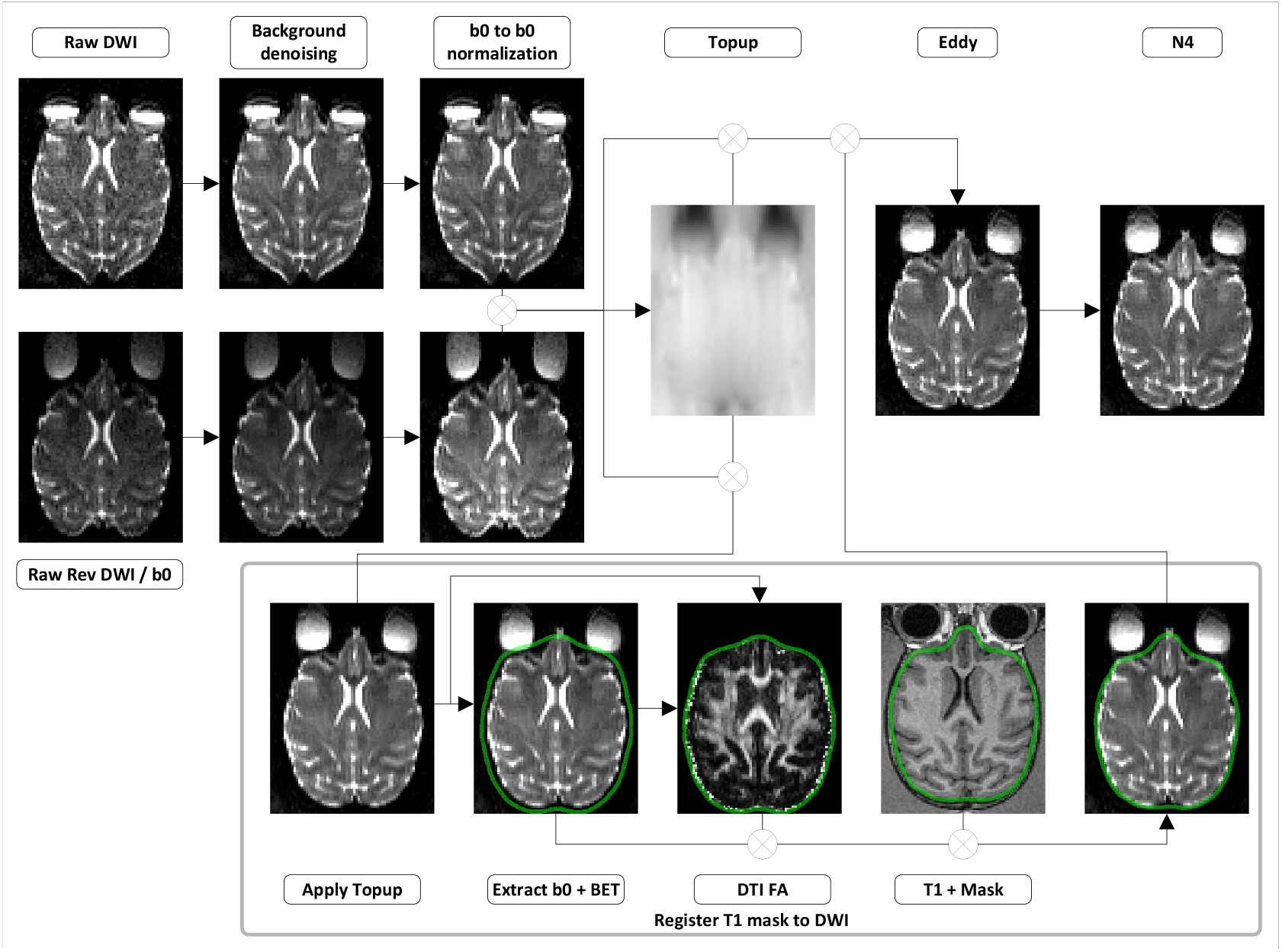
DWI preprocessing steps. (Background denoising) Input 4D DWI from both phase encoding directions are denoised to remove background noise contamination. (b0 to b0 normalization) The signal is normalized so all b0 volumes in both phase encoding directions exhibit the same mean value. Each 3D diffusion volume is normalized using as reference either the b0 volume coming before it, after it or a linear interpolation of both. (Topup) b0 volumes from both phase encoding directions are concatenated and a deformation field is estimated. (Apply Topup) This field is applied on the forward phase encoded DWI to produce an undistorted 4D DWI. (Extract b0 + BET) From the undistorted 4D DWI is extracted a mean b0 volume, upon which is computed a brain mask. All voxels outside the brain mask are set to 0 in the mean b0 volume and the undistorted 4D DWI. (DTI FA) The masked undistorted 4D DWI is used to compute a DTI model and obtain a FA volume. (Register T1 to DWI) A transformation is computed by registering the T1 volume to both the mean b0 and the FA volumes. (Apply Transform to Mask) The T1 mask is transformed to the DWI space using the computed transformation. (Eddy) Motion and eddy currents correction is computed using the deformation field from Topup, the T1 mask in DWI space and the 4D DWI from both phase encoding directions. (N4) The 4D DWI is corrected for intensity distortions.

**Figure 3.**
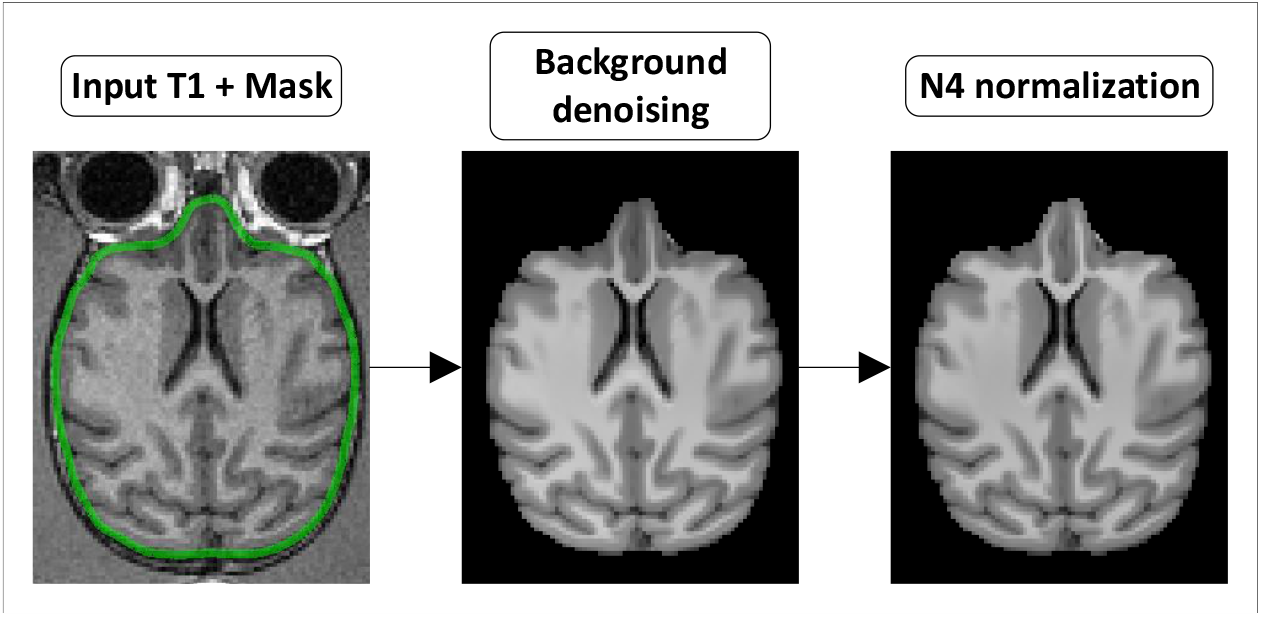
T1 preprocessing steps. (Background denoising) The T1 images is denoised to remove background noise contamination. (N4 normalization) The denoised T1 image is then corrected for intensity distortions.

**Figure 4.**
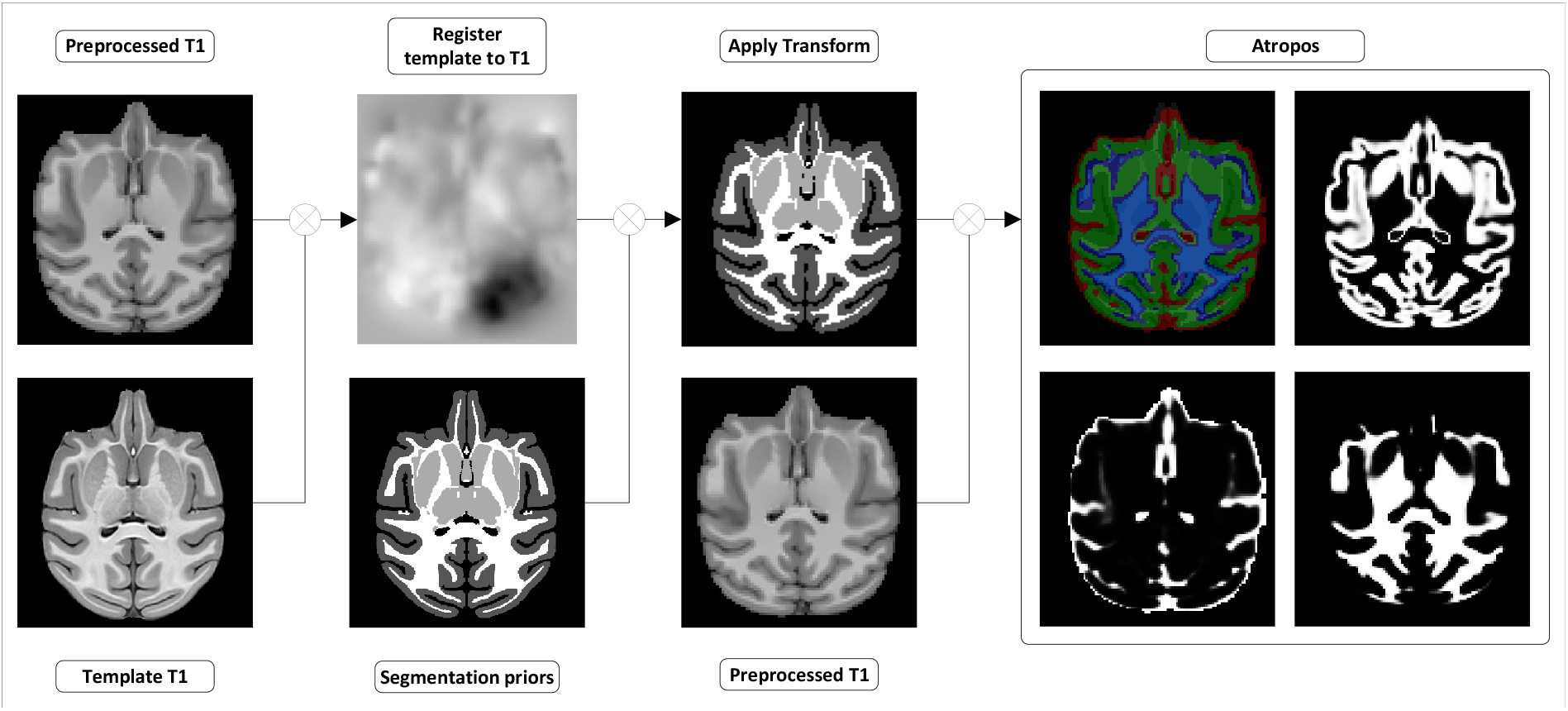
Tissues and white matter segmentation steps. (Register template to T1) A transformation from the template T1 to the preprocessed T1 is computed. (Apply transform) The transformation is used to move the segmentation prior into the preprocessed T1 space. (Atropos) Ants Atropos is run, with the transformed segmentation priors, to compute from the preprocessed T1 segmentation masks and partial volume fraction maps of white matter (WM), grey matter (GM) and cerebro-spical fuild (CSF).

All other scopes are using Nextflow workflow objects and only specify the flow of data between processes and other workflows they include. The input scope gathers pieces of code necessary to translate the internal representation in disk memory of the input data into a data structure that is digestible by a pipeline. When included, it enforces the input convention of its pipeline. It can be changed to fit a specific study or project. The pipeline scope contains all complete pipelines. A valid pipeline uses one or more input convention from the input scope and describes as minimally as possible the flow of data between processes and sub-workflows it includes. All data flow descriptions that can potentially be reused are transferred to the workflow scope. This scope contains sub-parts of the processing that form a coherent ensemble, yet cannot be specified by a single process. The workflows in this scope uses processes and other workflows as building blocks and define the passage of data between them to form a processing sequence. An example of such workflow would be the execution of Topup (a light purple workflow in figure 1), including data preparation, which executes the following processes sequentially : selection of b0 volumes from the different phase encoded DWI, concatenation of b0 volumes, computation of Topup distortion field, application of the field to the DWI.

### 2.2 Processing pipeline

#### 2.2.1 Data input

In addition to the DWI volume and its associated bval/bvec files describing the gradient sampling, the pipeline requires as input a T1 anatomical image. The user can also supply masks in either or both diffusion and anatomical space, as well as partial volume maps of white matter (WM), grey matter (GM) and cerebrospinal fluid (CSF), in anatomical space. Reverse phase acquired diffusion images can be provided, as either a single b0 volume, multiple b0 organized as a 4D volume or a full DWI 4D volume with bval/bvec files. For some of the processing steps, the pipeline also requires the input of some metadata associated to the acquisition of the DWI volumes. Metadata parameters consistent across all images can be supplied using the Nextflow configuration file. In case of varying parameters between the images, a json file must be provided alongside both forward and reverse acquired DWI volumes.

#### 2.2.2 Processing steps

The following section presents all currently implemented steps in our pipeline, which are also available in our processes and workflows library. Even though we highly recommend using them all in the order prescribed to process raw anatomical and diffusion weighted images, their execution is by default optional and can be individually turned on and off using the Nextflow configuration file, for instance to allow the computation of the models and measures on already preprocessed data or to shorten the execution time by turning off steps deemed unnecessary after input data quality control.

##### 2.2.2.1 Preprocessing

###### Background noise

dMRI images are affected with background noise following either a rician [Gudbjartsson and Patz (1995)] or a chi-squared distribution [Luisier et al. (2012)] and more often than not present quite poor signal-to-noise ratio. Some patch based algorithms [Manjón et al. (2008)] enable correction of such types of noise, but require long execution time and vast amounts of memory space, which can be cumbersome given the large size of diffusion volumes. A good trade-off is to use a Principal Component Analysis to extract new dimensions isolating the different sources of noise from the data. This technique has proven its abilities at enhancing the quality of diffusion data, even if it does not take into account the true nature of the noise and only corrects for normally distributed distortions. We thus run dwidenoise [Cordero-Grande et al. (2019)][Veraart et al. (2016b)][Veraart et al. (2016a)], a MP-PCA denoising algorithm available in Mrtrix3 [Tournier et al. (2019)], at the beginning of the preprocessing tree for diffusion, on forward and reverse phase acquired diffusion volumes, and Non-Local means [Manjón et al. (2008)] denoising on reverse phase acquired b0 volumes. Novel techniques using self-supervised learning such as Patch2Self [Fadnavis et al. (2020)] and deep convolutional neural networks [Kawamura et al. (2021)] could also easily be integrated in our modular framework, but have been discarded for the current implementation, since they still require to be thoroughly validated.

###### b0 to b0 normalization

High spatial and angular resolution diffusion images require long acquisition times. This causes heating in the acquisition coils, translated into a drift of signal intensities over time [Vos et al. (2017)]. To compensate for this effect, each pair of forward and reverse acquired images are normalized to the mean value of the first group of b0 volumes found in the forward image. To better correct the diffusion weighted volumes, a linear combination of the means of the groups of b0 volumes found before and after it is used. This behavior can be modified to accommodate datasets with different deviations between b0 volumes and diffusion images, such as taking only the b0 before or after a group of diffusion directions to compensate for the signal drift.

###### Gibbs ringing

Since the spatial MRI images are reconstructed by inverse Fourier Transform from a subset only of points sampled through k-space, images can suffer from wave-like artifacts at the interface between different tissues [Czervionke et al. (1988)]. When uncorrected, this leads to biases in models and metrics. We thus included Gibbs ringing correction as a step just after background denoising, using mrdegibbs [Kellner et al. (2016)] from Mrtrix3 [Tournier et al. (2019)]. However, since most images are acquired using partial fourier and that this algorithm does not support it, this step is skipped by default.

###### Susceptibility

To correct for those artifacts, we use Topup [Andersson et al. (2003)] from the FSL [Smith et al. (2004)] library. Topup is run on the extracted b0 of forward and reverse phase acquisitions to correct for susceptibility induced distortions. The extraction of b0 can be configured to :

- Take the first b0 appearing in a dwi image
- Take all the b0 in the dwi image
- Take the averages of b0 in the dwi image
- Take averages of continuous series of b0 in the dwi image

Since Topup is not parallelized, the pipeline comes initially configured to extract the average b0 for this preprocessing step. For this study, this configuration has proven to provide a good approximation of the distortion field while lowering substantially the execution time of the algorithm. However, in the case of variations in the susceptibility field across the diffusion volumes, sampling more than one b0 per image is recommended.

###### Brain masking

Since most of the subsequent steps in the pipeline require a mask for better and/or faster computation, we compute one using FSL bet [Smith (2002)] on the mean b0 volume, using the one corrected by Topup if it was run. However, the execution of bet on monkey images has proven to be unstable and often leads to bleeding of the computed mask into skull and background regions. To circumvent this problem, we highly recommend to pass as input data either a mask aligned to the diffusion or computed on the T1 anatomical image, that has been quality checked and manually fixed. In the latter case, the b0 bet mask will be used only to help register the T1 to the mean b0 image + FA (via ANTs [Avants et al. (2008)] using a combination of rigid, affine and non-linear transformations) in order to transform the T1 mask to diffusion space.

###### Motion and Eddy currents

Eddy currents are naturally occuring in tissues imaged at the MRI due to the rapid variations of the imaging gradients [Ahn and Cho (1991)]. They cause local distortions in the magnetic field perceived by the protons spins, leading to spatial distortions in the acquired images. To correct for those, we use FSL Eddy [Andersson et al. (2018)][Andersson et al. (2017)][Andersson and Sotiropoulos (2016)][Andersson et al. (2016)], which also corrects for patient motion. This step is parallelized both on CPU (using openMP) and GPU (using CUDA on a Nvidia graphics card). When using the latter, execution is more efficient and one can also perform outlier detection and replacement, as well as slice-wise corrections.

###### Intensity distortions

MRI equipment - gradients and reception coils - are not perfectly accurate and images can suffer from variation in intensities that are not related to the diffusion process of the acquired tissues. This leads to distortions in the diffusion measures recovered from the models fitted to the signal. This effect can also be caused by the presence of high susceptibility materials near or inside the imaged brain. To correct for those, we run N4 [Tustison et al. (2010)] intensity normalization from ANTs on the mean b0 of each diffusion. We then apply the computed intensity transformation to all diffusion directions acquired.

###### Background noise

Since the T1 image is composed of a single 3D volume, Non-Local Means Denoising [Manjoń et al. (2008)] becomes tractable in a reasonable amount of time. It is run using a rician noise model and parallelized over multiple CPU cores, via the algorithm available in Dipy [Garyfallidis et al. (2014)]. This steps has proven to improve segmentation and registration [Theaud et al. (2020)][Constanzo et al. (2018)] and has thus been integrated as an optional step in the pipeline.

###### Intensity distortions

As for diffusion, the T1 is corrected for local intensity distortions caused by the MRI equipment using ANTs N4 [Tustison et al. (2010)] intensity normalization algorithm.

##### 2.2.2.2 Upsampling

Once denoised, both T1 and the diffusion volumes are upsampled to a finer resolution defined in the Nextflow configuration file using Dipy [Garyfallidis et al. (2014)]. This step enhances anatomical features present in the diffusion volumes, brings both T1 and diffusion on a common spatial grid, allowing the use of masks and segmentations subsequently computed on the T1 in reconstruction and measures algorithms acting on the diffusion, and is also known to improve the results obtained from tractography [Dyrby et al. (2014)].

##### 2.2.2.3 Segmentation

###### Tissue segmentation

Segmentation is performed on the T1 using ANTs Atropos [Avants et al. (2011)]. The default configuration of the pipeline uses AFNI NMT v2.0 segmentation [Jung et al. (2021)] as priors, after registering the NMT template to the T1 image, using ANTs [Avants et al. (2008)] (rigid + affine + non-linear, MultiLabel interpolation). Tissue masks are also extracted from the partial volume fractions outputed by Atropos by thresholding. A safe white matter mask, exempt of partial volume effects with both CSF and gray matter, is computed for future usage such as limiting the number of possible voxels used to compute the single fiber response necessary for constrained spherical deconvolution.

###### White matter parcellation

The default white matter atlas available in the pipeline is the UWDTI atlas [Adluru et al. (2012)]. It provides templates in the form of DTI measures, of which we chose the FA to perform registration. The FA is calculated from a DTI fit on the upsampled DWI data. Then a transformation from template space to subject space is computed using ANTs [Avants et al. (2008)] (rigid + affine + non-linear) and applied to the atlas using MultiLabel interpolation.

##### 2.2.2.4 Registration of T1 to DWI space

Once segmentation is computed, the upsampled T1 image is registered to the upsampled DWI space using a combination of the mean b0 volume and the FA obtained from a DTI fit. The registration is computed using ANTs and a sequence of Rigid, Affine and Non-Linear transformations. This transformation is then applied to all segmentation masks using Nearest Neighbour interpolation, to label maps resulting from tissue and white matter segmentation using MultiLabel interpolation and to partial volume maps from tissue segmentation with Linear interpolation.

##### 2.2.2.5 Reconstruction

The pipeline offers 3 different models for the reconstruction of the diffusion process. To enable its usage on low angular resolution data or data sampled across few shells at low b-values, the single diffusion tensor model [LeBihan et al. (2001)] (DTI) can be fitted. It requires a minimum of 6 directions, though a uniform sampling of gradients over the sphere is preferable. To limit the effects of exchange and restriction occurring at high b-value, which this model cannot represent, only shells up to b = 1300*mm/s*^2^ are used [Jones and Basser (2004)]. This behavior can be modified by the user if need be. For data acquired on a greater number of shells, with higher b-values, such as HARDI [Hosey et al. (2005)][Tuch et al. (2002)] and CUSP[Scherrer and Warfield (2012)], two higher-order models are available : the fiber orientation distribution function (fODF) through Constrained Spherical Deconvolution [Tournier et al. (2007)] (CSD) and the DIstribution of Anisotropic MicrOstructural eNvironments in Diffusion-compartment [Scherrer et al. (2016)] (DIAMOND), a multivariate gamma distribution over tensors.

##### 2.2.2.6 Measurement

###### DTI fit

Diffusion tensors are computed using Dipy and a Weighted Least Square Fit. In addition to the tensors, eigenvalues and eigenvectors, it outputs axial (*λ*_1_) and radial diffusivity (*µ*(*λ*_2_, *λ*_3_)), fractional and geodesic anisotropy, the mode and norm of diffusion tensors. Figure 5 presents some of those metrics. It also allows the output of validation maps, such as the residuals of the fit, the standard deviation across diffusion volumes displaying pulsation and misalignment artifacts, and a map of physically implausible voxels, representing the areas where the diffusion weighted signal presents higher values than the b0 mean.

**Figure 5.**
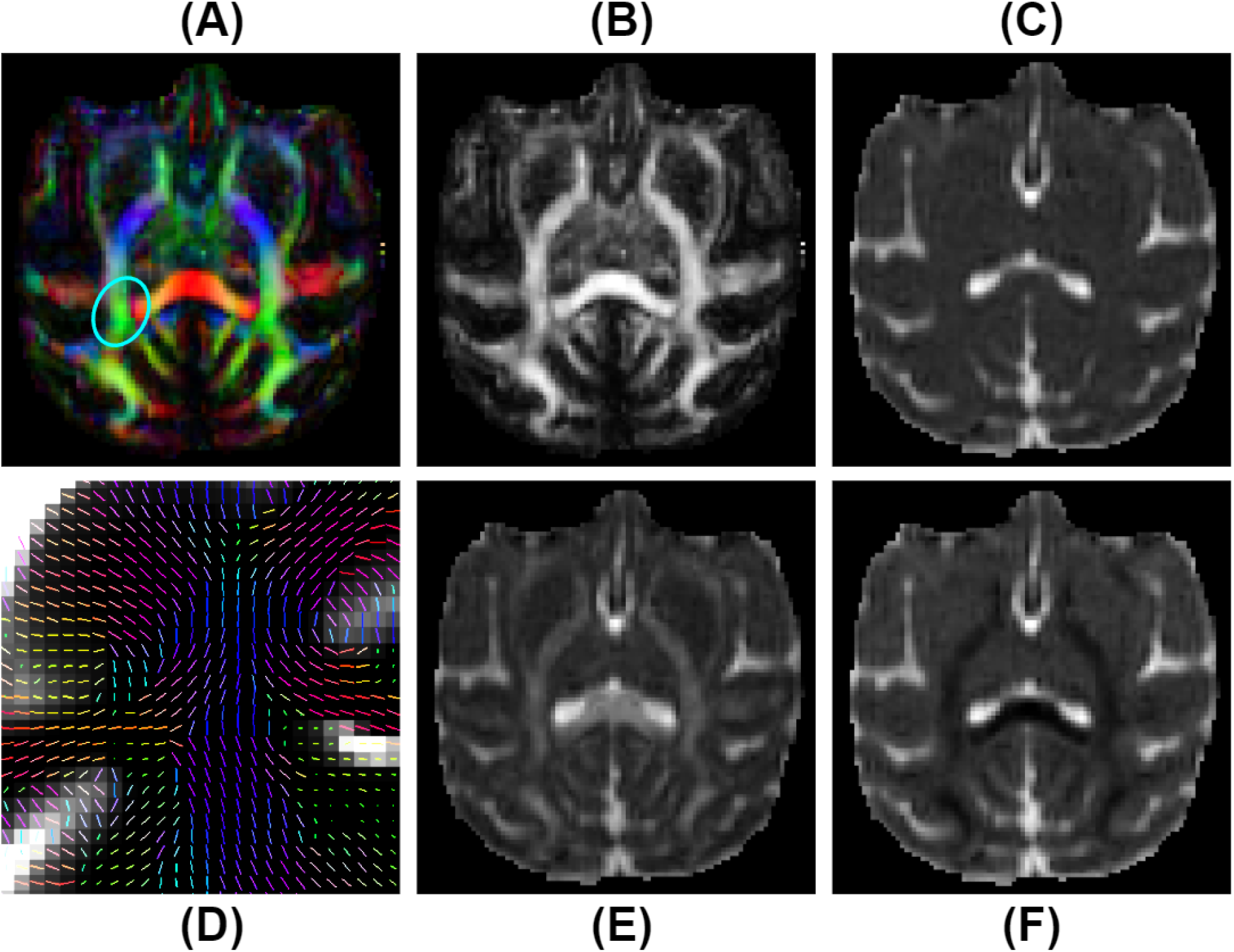
DTI measurements on a selected subject from the Aix-Marseille site. (A) RGB map, (B) fractional anisotropy (FA), (C) mean diffusivity (MD), F) radial diffusivity (RD), (E) axial diffusivity (AD) and (D) the principal direction of the estimated tensor in the centrum semi-ovale (region circled in blue in (A)).

###### CSD fit

The response function and spherical deconvolution fit are also computed using Dipy. If a tissue segmentation is available at this step in the pipeline and the input DWI contains multiple b-value shells, a Multi-Shell Multi-Tissue approach is used [Jeurissen et al. (2014)]. Otherwise, the pipeline reverts to using a Single-Shell Single-Tissue algorithm [Tournier et al. (2007)]. The response is obtained using a subset of single-fiber white matter voxels (200 per default) which are presenting a high enough FA value (between 0.55 and 0.75 per default) and reside inside the safe white matter mask priorly computed. The peaks are then extracted from the fitted fODF, selecting only the ones presenting an amplitude greater than 1.5 times the maximal amplitude in isotropic voxels [Dell’Acqua et al. (2012)] (figure 6 : F). The latter are obtained using a threshold on mean diffusivity (a minimum of 2.6*e*^*™*3^*mm*^2^) and fractional anisotropy (0.15). Those threshold are also used to compute the Number of Fibers Orientations and the maps of voxel-based total (first SH coefficient), sum (integral on the sphere) and maximal Apparent Fiber Density [Dell’Acqua et al. (2012)][Raffelt et al. (2012)], as well as the RGB mapping of orientations on the sphere (figure 6 : A-E).

**Figure 6.**
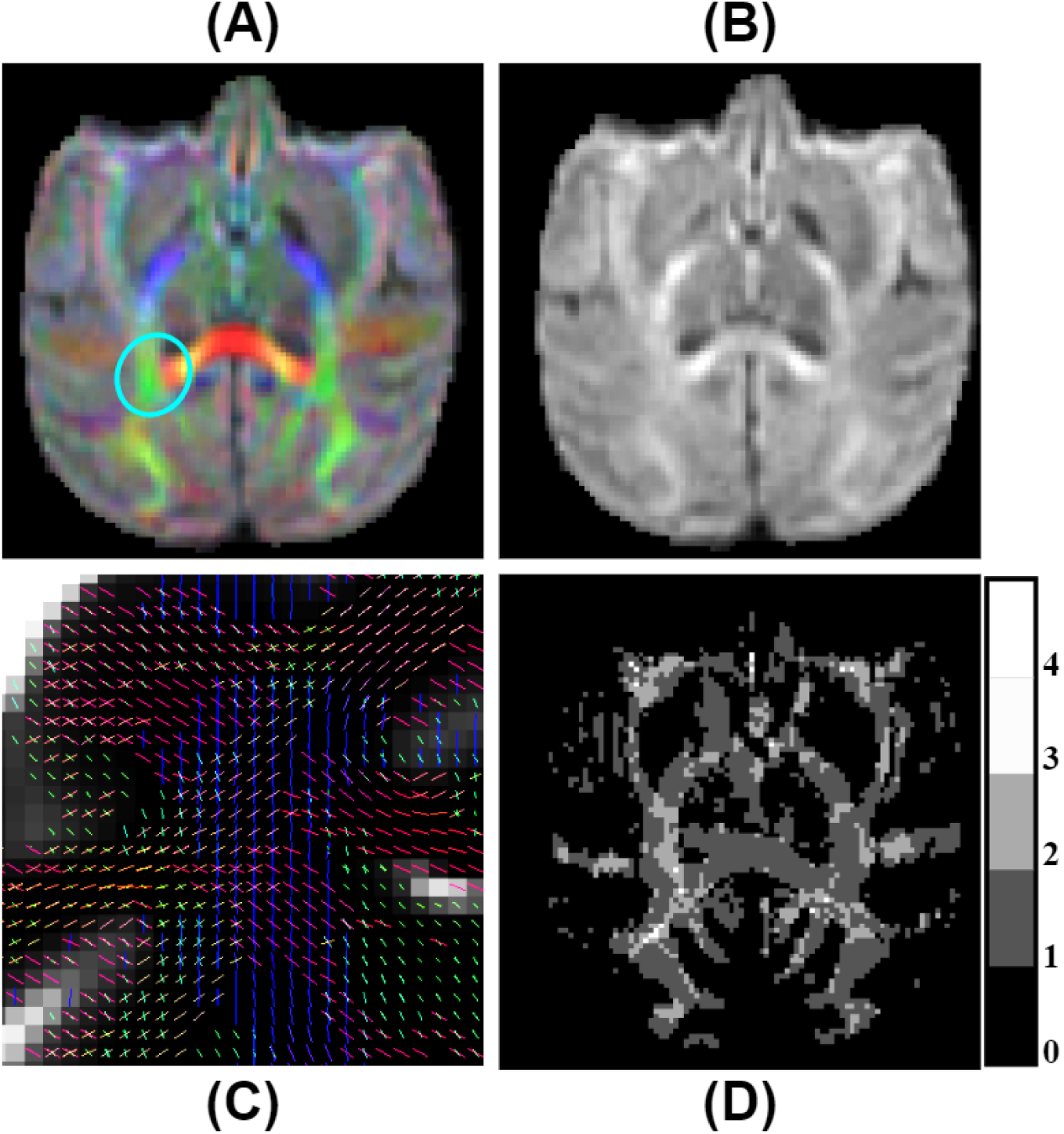
fODF measurements on a selected subject from the Aix-Marseille site. (A) RGB map, (B) total apparent fiber density (AFDt), (D) number of fiber orientations (NuFO) and (C) fODF peaks in the centrum semi-ovale (region circled in blue in (A)).

###### Multivariate Gamma Distribution Tensor Fit

The Multi-Tensor fit is acquired using Diamond [Scherrer et al. (2016)], configured to allow for a maximum of 2 tensors (fascicles) per voxel. As default, the fit also outputs the fraction of free-water. Classical tensor measures (FA, AD, RD, MD, RGB) are estimated on each fascicle separately and are used to compute their mean and max counterparts. In addition is generated a map of the fraction of each fascicle per voxel and of their main peaks, and other statistical quantities such as the mean and standard deviation of isotropic and anisotropic diffusivities [Reymbaut and Descoteaux (2019)]. A subset of metrics is displayed in figure 7.

**Figure 7.**
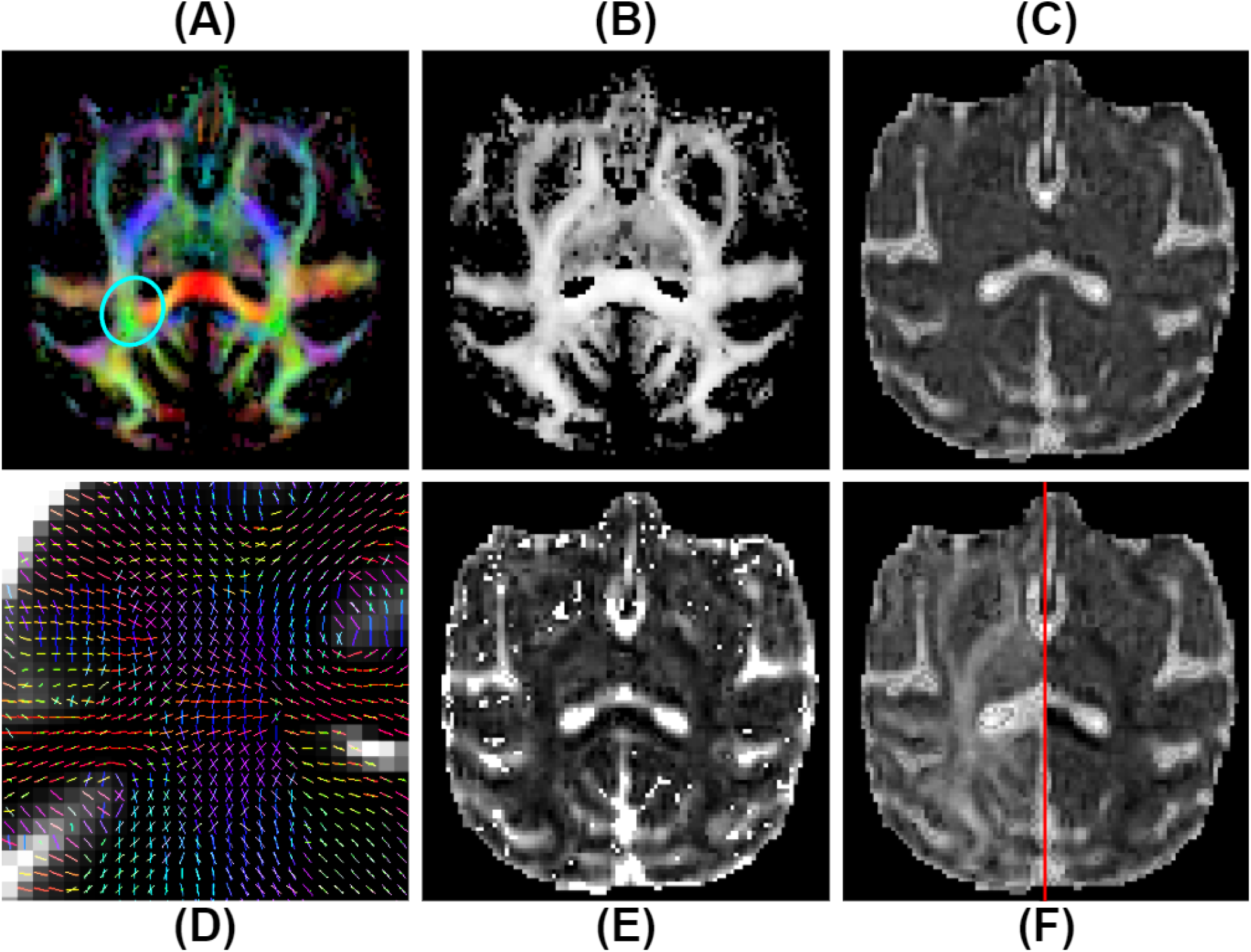
Multi-tensor measurements on a selected subject from the Aix-Marseille site. (A) RGB map, (B) maximum fascicle FA (max fFA), (C) mean diffusivity (MD), (F) axial diffusivity (AD) on the left, radial diffusivity (RD) on the right, (E) free water fraction (fFW) and (D) principal direction of the estimated mean tensor of each fascicle in the centrum semi-ovale (region circled in blue in (A)).

### 2.3 Pipeline reproducibility

To evaluate correctly the potential variations in metrics between the several subjects of the PRIME-DE databases, it is a requirement that the processing carried on the data be as reproducible as possible. To do so, all algorithms used by the pipeline that require the execution of a random process have seen their seed enforced to the same number across all subjects and sites. To evaluate the reproducibility, we did a test-test analysis, using the 4 subjects from the aix-marseille database, on which the pipeline was run 3 separate times, using the same configuration and singularity image. The image intraclass correlation coefficient (I2C2) [Shou et al. (2013)] was computed using equation 1 on relevant DTI, fODF and multi-tensor metrics.

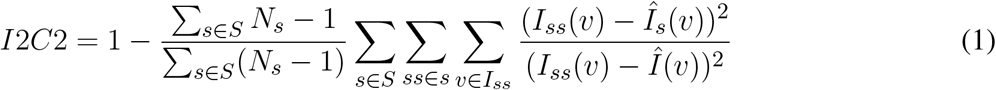

Here, *I*_*ss*_ refers to a modality, mask or measurement to evaluate for a specific session *ss* belonging to a subject *s, Î*_*s*_ is the mean image for a specific subject *s* taken over all its sessions and *Î* is the mean image taken over all sessions and all subjects.

### 2.4 Study design and data selection

This study focuses at evaluating variability in diffusion weighted (DW) measurements across the different sites available in the Primate Data Exchange (PRIME-DE) project, as well as between the MRI scanner vendors and models used by them to acquire the images. From the 25 sites available in the database, 8 of them were identified as containing diffusion data, and only 4 with a fair number of subjects (*N* ≥ 4) : Aix-Marseille University (4 subjects, Siemens Prisma scanner), University of California, Davis (19 subjects, Siemens Skyra scanner), Mount Sinai School of Medicine - Philips (9 subjects, Philips Achieva scanner) and Mount Sinai School of Medicine - Siemens (6 subjects, Siemens Skyra scanner). All subjects are Macaca Mulatta primates, anesthetized before acquisition. The scan sequences parameters can be found in table 1. All datasets were acquired on 3T systems, with DWI at spatial resolutions from 0.7 to 1.4 *mm*^3^, with either isotropic or rectangular voxels, and anatomical images at voxels sizes ranging from 0.3 *mm*^3^ to 0.8 *mm*^3^ isotropic.

**Table 1.** Acquisition parameters for the sites of the PRIME-DE database containing diffusion data.

| SITE | SCANNER | # subjects | T1 | DWI |  |  |  |
| --- | --- | --- | --- | --- | --- | --- | --- |
|  |  |  | voxel size | voxel size | b-values | # directions | # b0 |
| Aix-Marseille | Siemens<br>Prisma 3T | 1 | 0.8mm<br>0.8mm<br>0.8mm | 1.25mm<br>1.25mm<br>1.25mm | 250<br>1000 | 6<br>64 | 5 |
|  |  | 1 | 0.8mm<br>0.8mm<br>0.8mm | 1mm<br>1mm<br>1mm | 250<br>1000 | 6<br>64 | 5 |
|  |  | 1 | 0.8mm<br>0.8mm<br>0.8mm | 1.25mm<br>1.25mm<br>1.25mm | 300<br>1000<br>2000 | 6<br>32<br>64 | 7 |
|  |  | 1 | 0.8mm<br>0.8mm<br>0.8mm | 1mm<br>1mm<br>1mm | 500<br>1000<br>2000 | 6<br>30<br>30 | 5 |
| UC-Davis | Siemens<br>Skyra 3T | 19 | 0.3mm<br>0.3mm<br>0.3mm | 0.7mm<br>0.7mm<br>1.4mm | 1600 | 60 | 6 |
| Sinai-Philips | Philips<br>Achieva 3T | 9 | 0.3mm<br>0.3mm<br>0.3mm | 1mm<br>1mm<br>1mm | 1000 | 120 | 2 |
| Sinai-Siemens | Siemens<br>Skyra 3T | 6 | 0.5mm<br>0.5mm<br>0.5mm | 1mm<br>1mm<br>1mm | 1000 | 80 | 10 |

### 2.5 Quality control

All input data was quality controlled on both DW and anatomical images. Images were controlled for a variety of possible artifacts, including motion, ghosting, aliasing, gibbs ringing, B0 and B1 field inhomogeneities and magnetic susceptibility. After evaluation, images from Mount Sinai School of Medicine - Siemens were excluded due to non-uniformity of contrasts across both DW and anatomical images from hyper-intensities caused by restraints positioned around the subject’s head. The 3 other sites presented high enough image quality and were judged good enough for the study.

### 2.6 Brain masking

Brain masks were computed prior to processing on the anatomical T1 images using DeepBet v1.0 [Wang et al. (2021)], a U-Net trained on macaque data from the PRIME-DE. All masks were quality controlled and manually fixed to prevent exclusion of brain voxels and inclusion of skull and eyes. The inclusion of this technology in the pipeline was considered, but was ultimately rejected due to the weight in memory of its dependencies, practically doubling the size of the container required to run the pipeline. Moreover, the lack of validation on databases other than the PRIME-DE, which was used to generate the segmentation models, and its insertion near the beginning of the processing chain was deemed too risky, since its failure at providing a good brain mask would impede on most of all subsequent steps executed.

### 2.7 Processing hardware

The processing was done launching the pipeline separately for all 3 selected databases to evaluate the effectiveness of execution. We used 3 nodes of the Compute Canada Beluga cluster, each equipped with 2 Intel Gold 6148 Skylake processors at 2.4 GHz clock speed, 186 Gb of RAM and 4 NVidia V100SXM2 graphics cards, 16 Gb VRAM.

### 2.8 Data quality evaluation

To quantify the increase in quality of the processed data, two measures, SNR and CNR, were computed inside each region of the registered tissue maps on the mean b0 volume of the diffusion weighted images. Topup correction and upsampling was performed on the raw data, to allow for the use of the tissue maps when computing their SNR and CNR. Our method thus does not quantify the effect of those steps on the amelioration of the data quality.

To calculate SNR, two different approaches were used to quantify the noise standard deviation on the b0. For DWI containing multiple b0 volumes, intermediate noise maps were extracted by subtracting each possible permutations of two b0. Those maps were then averaged to obtain a mean noise map, which was used to compute the noise standard deviation in each region. SNR was then calculated using equation 2 [Narasimhan and Jacobs (2002)]. For DWI with a single b0 volume, a background mask was extracted using the technique presented in [Balan et al. (2012)] and the noise standard deviation was calculated from the voxels of the b0 inside it. SNR was calculated using equation 3. In both equation 2 and 3, *b*0 refers to the b0 image, *N* to the extracted mean noise map, *T*_*mask*_ to the mask of the tissue of interest and *B*_*mask*_ to the image background mask. *µ*_*M*_ (*I*) and 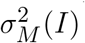 (*I*) respectively refer to the mean and variance of all voxels of an image *I* included in a mask *M* .

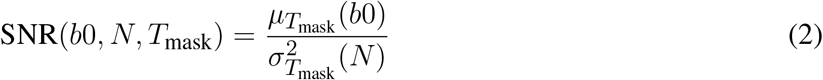

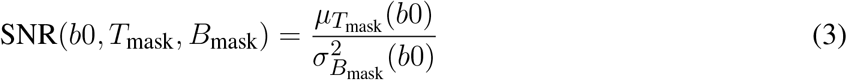

CNR was computed on the b0 by comparing white matter with gray matter regions using equation 4.

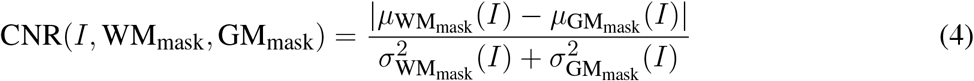

We also performed a variability study in tissue maps (WM and GM) of several metrics extracted from the diffusion models available in the pipeline. Using the average over sessions of subjects, we analyzed inter-subject variability separately for each database (equation 5), intra-vendor variability for Siemens scanners (UC-Davis and Aix-Marseille) (equation 8) and inter-vendor variability using all sites (equation 7). Since UC-Davis and Sinai-Philips presented multiple time points per subject, intra-subject variability was also computed (equation 6).

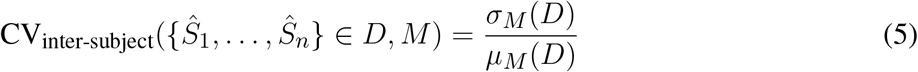

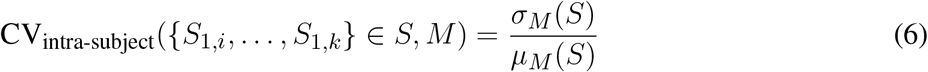

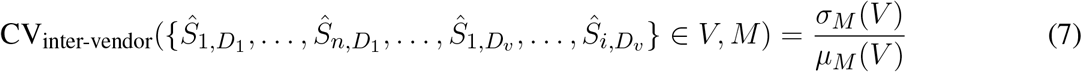

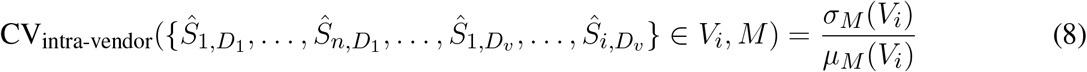

Here, *S* denotes a set of images (T1, DWI, masks and measures) of a single subject, across all of its repetitions, *D* a single acquisition site (Aix-Marseille, Sinai-Philips or UC-Davis), *V* a vendor (Philips, Siemens) and *M* a mask defining a region of interest in which is evaluated the variation measure.

## 3 RESULTS

### 3.1 Processing time

Per process execution time is displayed in figure 8 separately for each database. Most of the processing time was occupied by 4 algorithms, namely Topup (∼66 min avg), Eddy (∼36 min avg), DIAMOND (∼31 min avg) and fODF metrics computation (∼15 min avg); those times are heavily bound to the diffusion image resolution, here ranging from 1mm to 1.25mm isotropic. Every other major step of the pipeline executed in about 5 minutes or less. It took about 6 hours processing the 4 dataset of the Aix-Marseille site, 25 hours for the 27 (9 subjects, 3 sessions) of Sinai-Philips and 34 hours for the 38 (19 subject, 2 sessions) of UC-Davis. Note that for UC-Davis, the number of iterations for Topup had to be increased, due to the intense susceptibility artifacts present in the images, thus drastically increasing the execution time.

**Figure 8.**
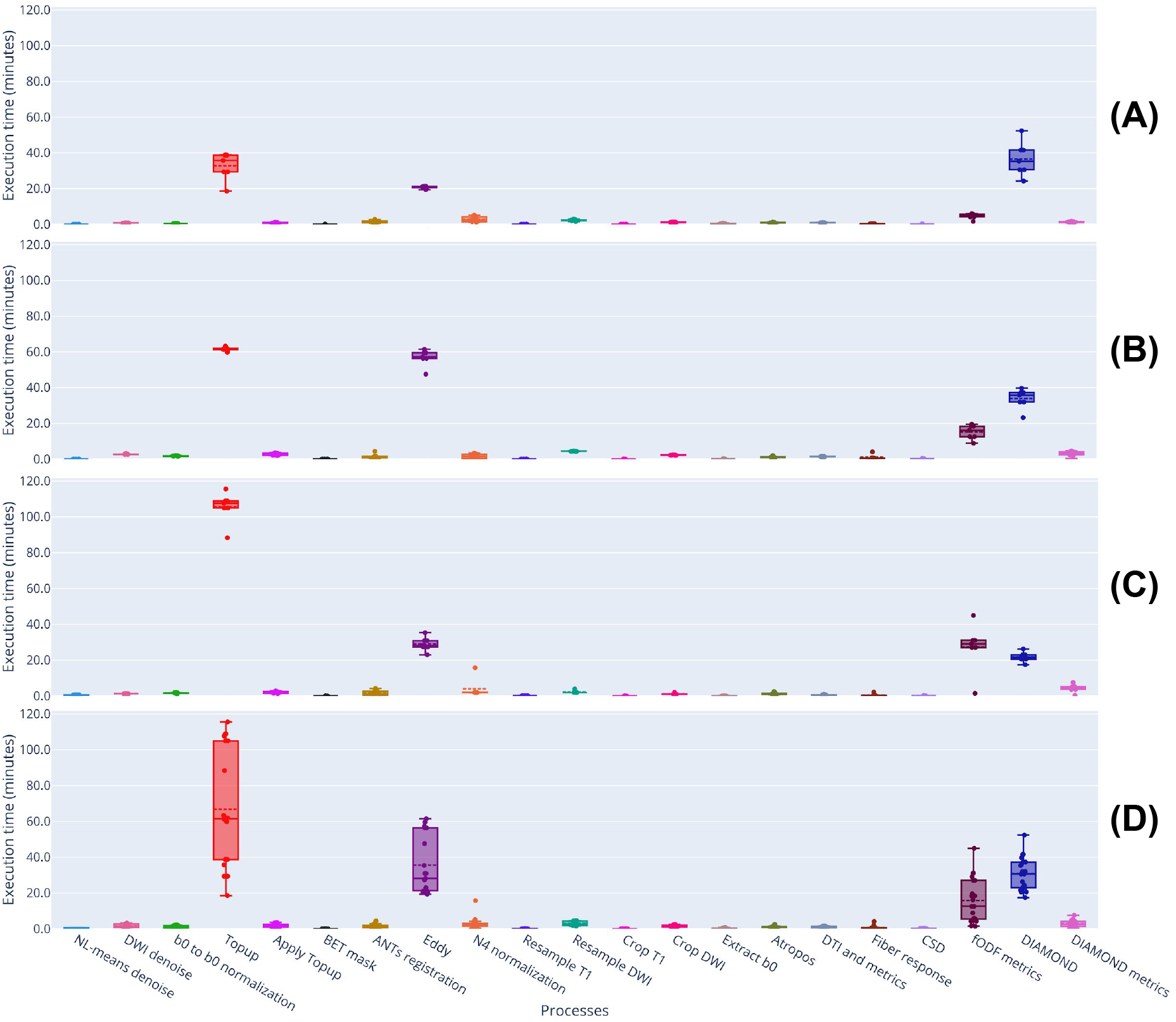
Execution time of processes run on datasets from the (A) Aix-Marseille, (B) Mount Sinai - Philips and (C) UC-Davis sites. (D) presents the global time consumption per process over all sites.

### 3.2 Image quality

Signal to noise ratio (SNR) is presented in figure 9, on the average b0 volume extracted from both raw and preprocessed DW images. With the exception of UC-Davis, the measure suggests an increase in image quality resulting from the different denoising algorithms used in the preprocessing part of the pipeline. For Sinai-Philips and UC-Davis, SNR distribution displays less variance after preprocessing, with a substantial increase of mean SNR for Sinai-Philips.

**Figure 9.**
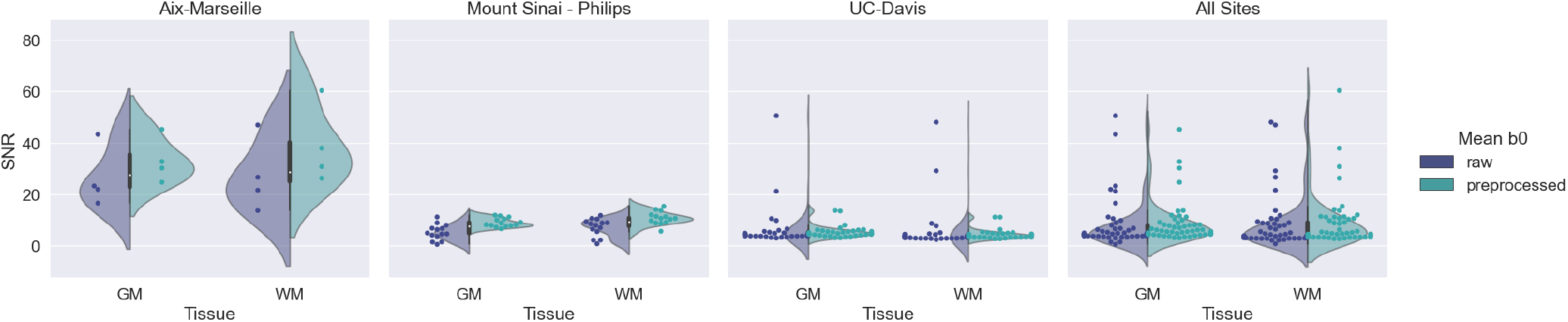
SNR of b0 images inside (WM) white matter, (GM) gray matter and (CSF) cerebro spinal-fluid tissues

Contrast to noise ratio (CNR) in white matter voxels is displayed in a similar fashion in figure 10. As for SNR, distributions of CNR display less variance after preprocessing for Sinai-Philips and UC-Davis. Mean CNR is also increased for Aix-Marseille and Sinai-Philips.

**Figure 10.**
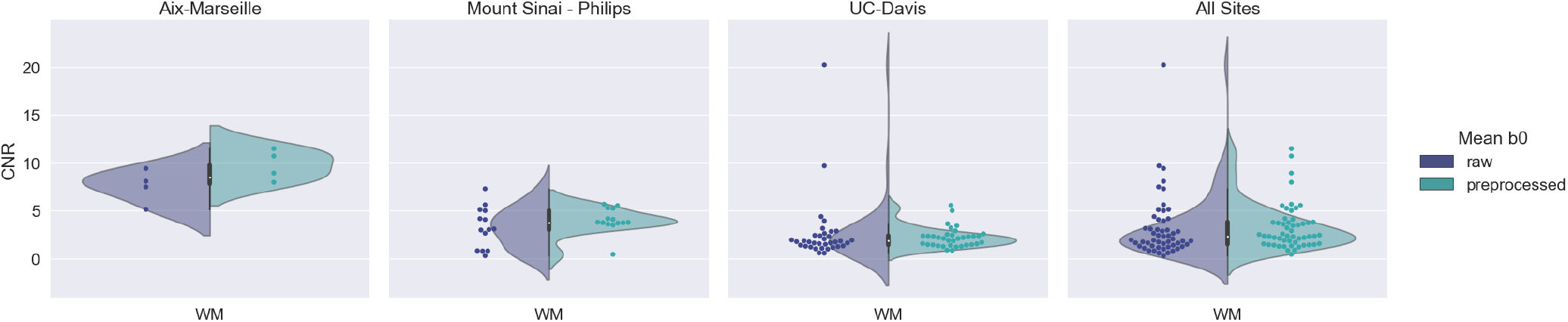
CNR of b0 images

### 3.3 Reproducibility

Table 2 shows reproducibility scores on revelant output images of the pipeline. For classical diffusion models (DTI and fODF), as well as for anatomical mask computing and registration processes, reproducibility is high, with I2C2 scores of 98% and more. Scores for the DIAMOND model averages around 97% for metrics computed over the fascicles (MD, AD, RD, FA, max fFA, free-water fraction) and 91% model selection, indicating that the outcomes of this algorithm are less reproducible. Gamma distribution parameters (kappa, kappaAD, hei and heiAD) do not perform as well, with kappa and hei scoring at 83% and heiAD and kappaAD at 52% and 63%.

**Table 2.** Reproducibility measures on images generated by the pipeline.

| Modality |  | Measure | I2C2 |
| --- | --- | --- | --- |
| Anatomy | T1 | WM PVF | 0.997 |
|  | T1 | GM PVF | 0.995 |
|  | T1 | CSF PVF | 0.990 |
|  | T1 | registration | 1.000 |
| Diffusion | DTI | MD | 0.991 |
|  | DTI | AD | 0.990 |
|  | DTI | RD | 0.991 |
|  | DTI | FA | 0.999 |
|  | fODF | AFD | 0.981 |
|  | fODF | AFD total | 0.997 |
|  | fODF | NuFO | 0.986 |
|  | DIAMOND | MD | 0.962 |
|  | DIAMOND | AD | 0.975 |
|  | DIAMOND | RD | 0.956 |
|  | DIAMOND | FA | 0.990 |
|  | DIAMOND | max fFA | 0.979 |
|  | DIAMOND | hei | 0.831 |
|  | DIAMOND | heiAD | 0.523 |
|  | DIAMOND | kappa | 0.831 |
|  | DIAMOND | kappaAD | 0.630 |
|  | DIAMOND | fascicle fractions | 0.968 |
|  | DIAMOND | free-water fraction | 0.981 |
|  | DIAMOND | model selection | 0.903 |

### 3.4 Variation of measurements

Figures 11 to 13 display the coefficient of variation of several metrics outputed by the pipeline, and figure 14 on the gamma distribution parameters estimated by the DIAMOND model, calculated in both white matter (WM) and gray matter (GM) masks.

**Figure 11.**
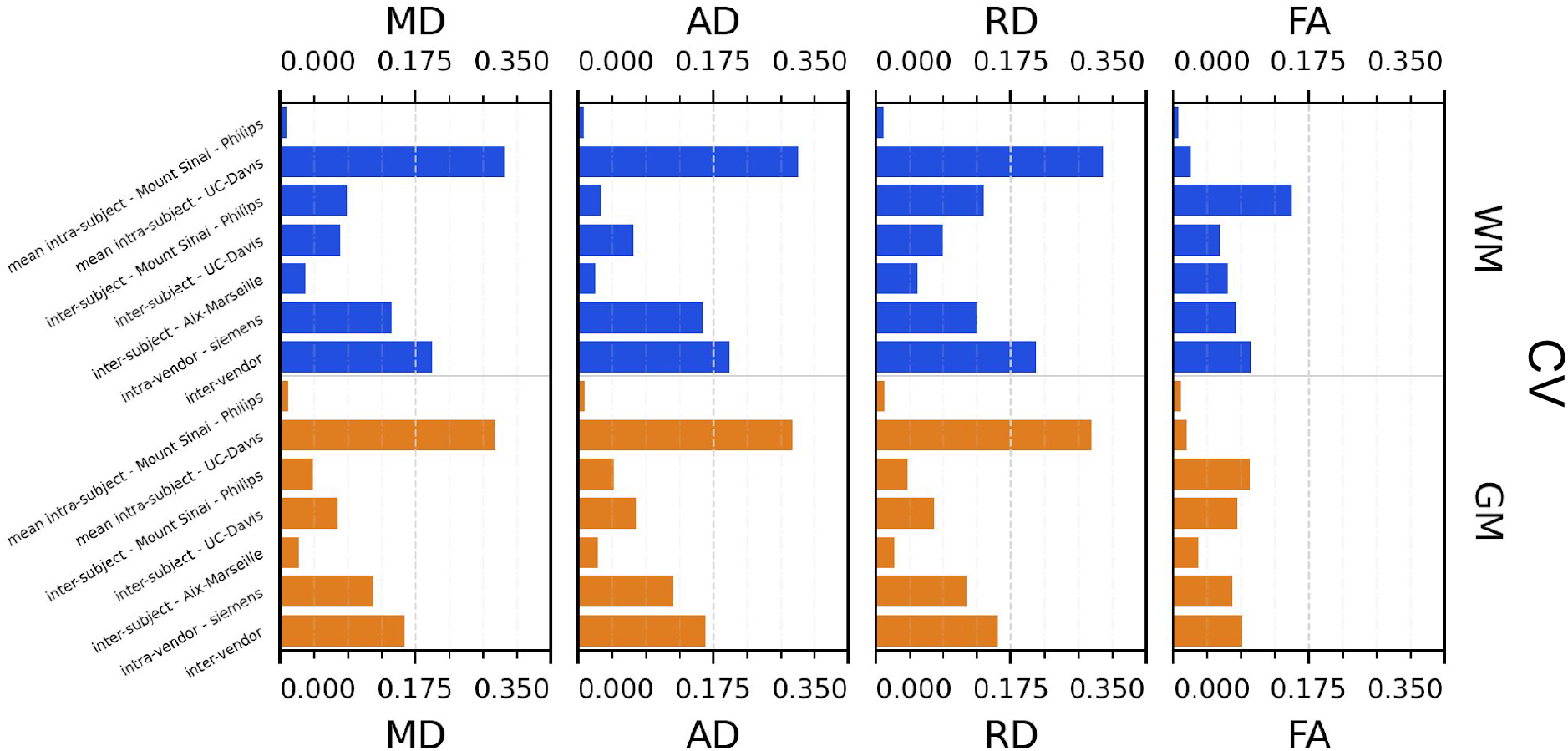
Coefficient of variation (CV) for DTI measurements : (MD) mean diffusivity, (AD) axial diffusivity, (RD) radial diffusivity, (FA) fractional anisotropy

**Figure 12.**
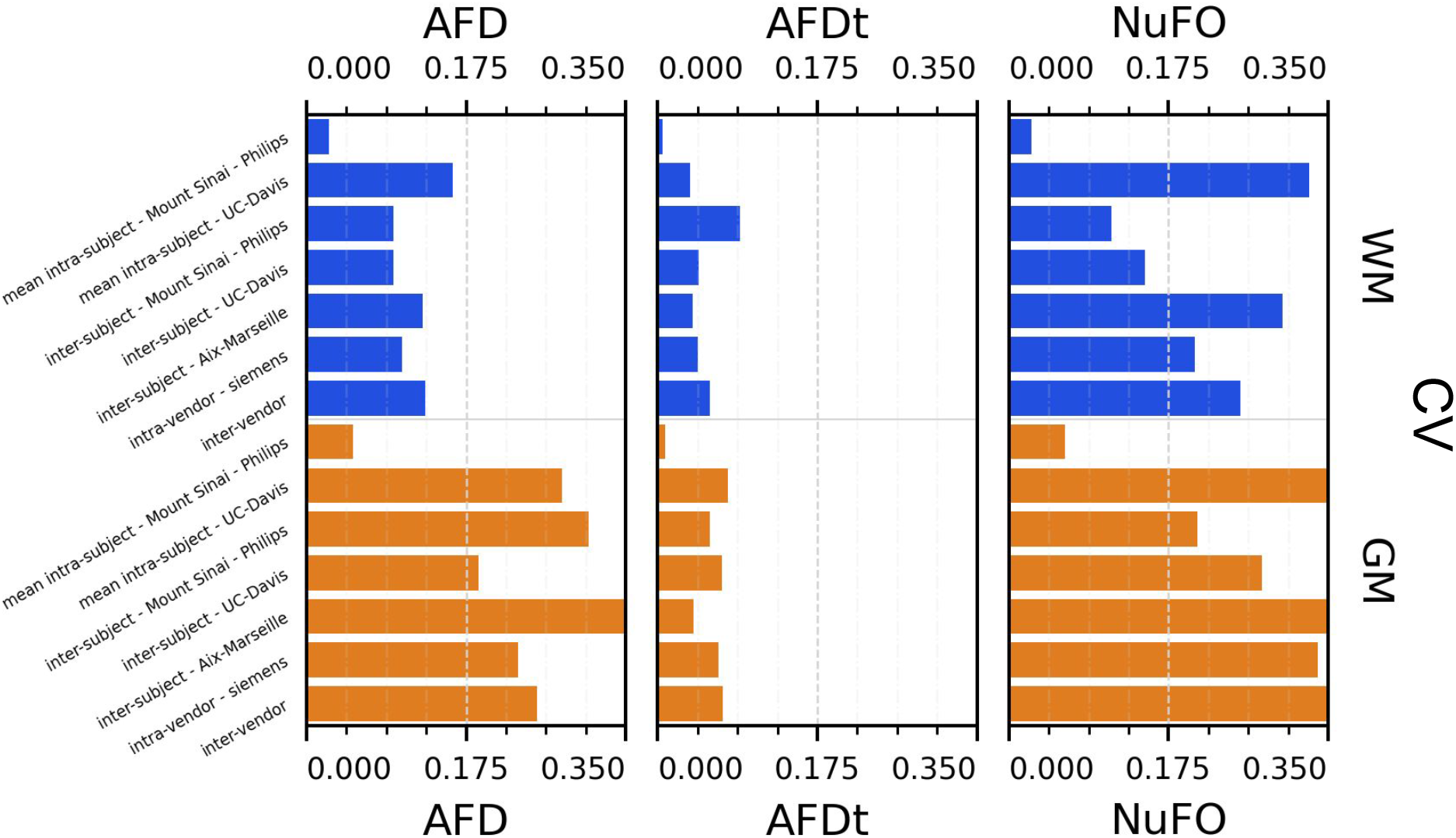
Coefficient of variation (CV) for fODF measurements : (AFD) max apparent fiber density, (AFDt) total apparent fiber density, (NuFO) number of fiber orientations

**Figure 13.**
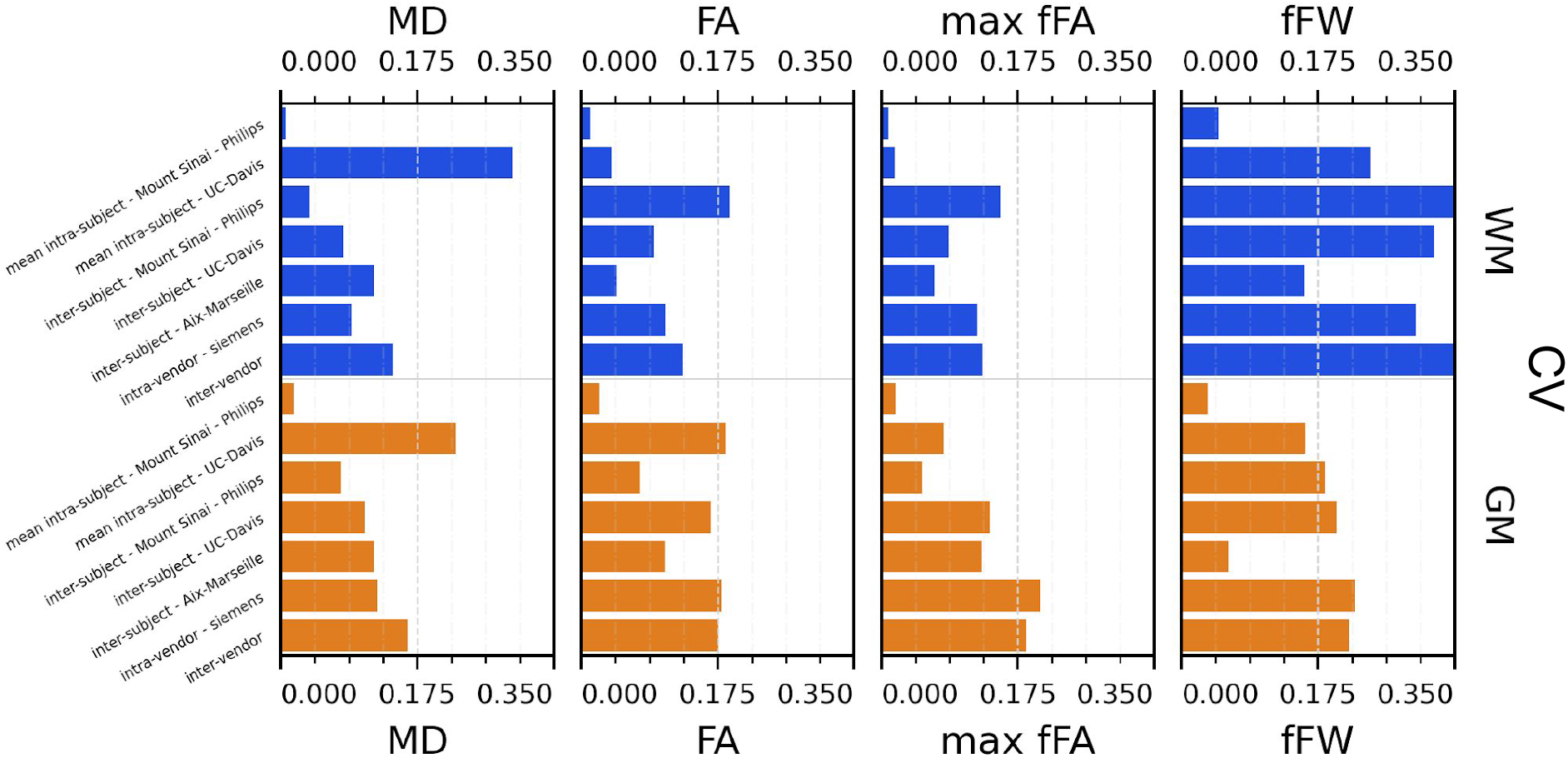
Coefficient of variation (CV) for diamond measurements : (MD) mean mean diffusivity among fascicles, (FA) mean fractional anisotropy among fascicles, (max fFa) maximum fascicle FA, (fFW) fraction of free-water

**Figure 14.**
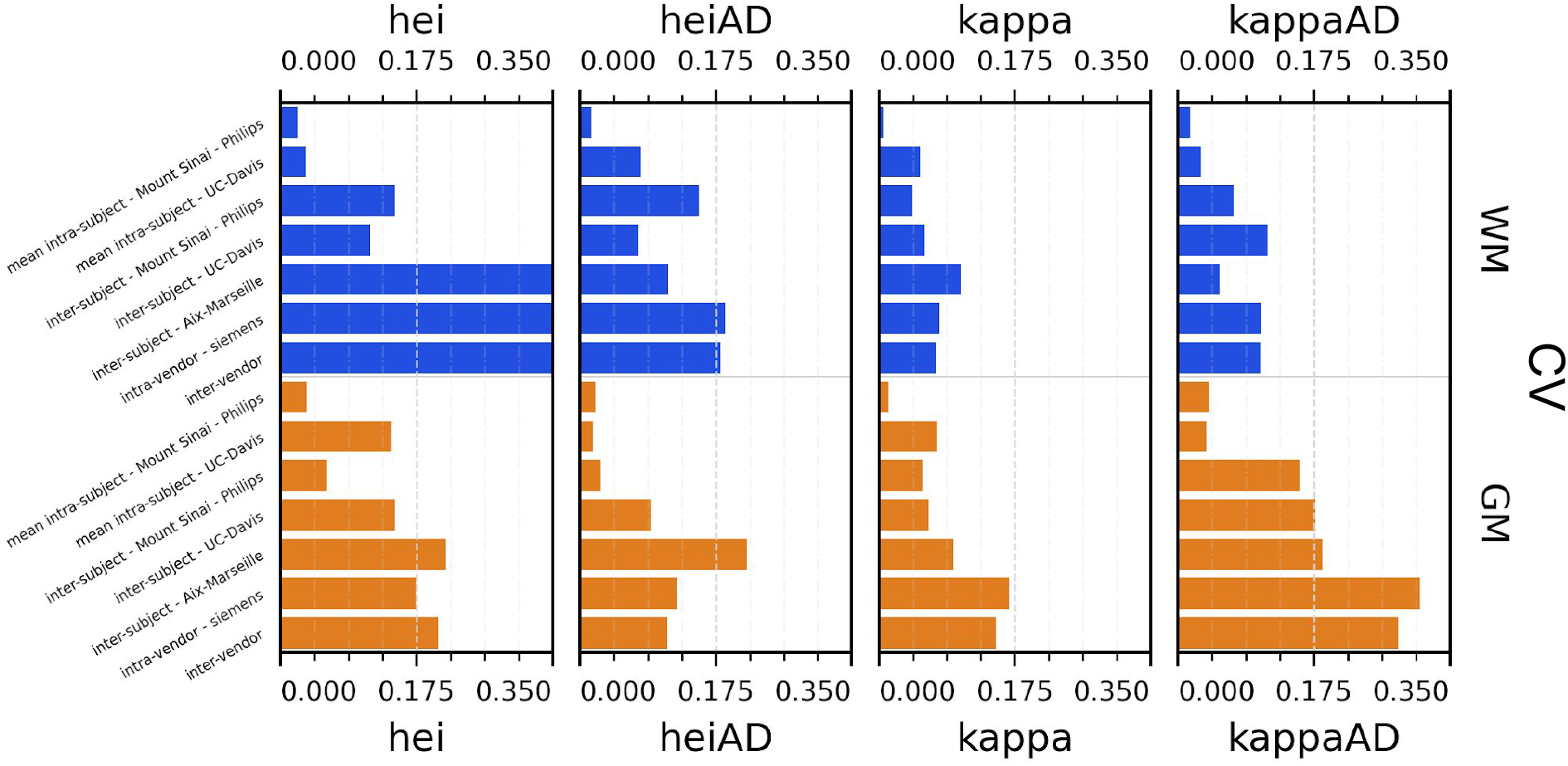
Coefficient of variation (CV) for diamond gamma distributions parameters : (hei) heterogeneity index, (heiAD) axial heterogeneity index, (kappa) kappa parameter of tensor distribution, (kappaAD) axial kappa parameter of tensor distribution

The Sinai-Philips site presents the least mean intra-subject variability overall at ≤2%, except for NuFO, AFD and free-water fraction (fFW), but the worst inter-subject variability in almost all metrics and parameters evaluated.

For DTI metrics, Aix-Marseille displays better inter-subject variability in both white matter (≤4% in MD and RD and ≤7% in RD and FA) and gray matter (∼3%) than Sinai-Philips. The portrait is reversed for fODF measurements, with the exception of AFDt, where the variability observed is greater (more than twice for NuFO in WM). For metrics computed on the DIAMOND model, the variability is better, with the exception of MD in WM and of all measures excluding free-water fraction (fFW) in GM. For gamma distribution parameters, variability is also higher, except for kappaAD and heiAD in WM.

UC-Davis exhibits good inter-subject variability in DTI measurements at less than 4% for MD, AD and RD and around 6% in WM and 8% in GM for FA. For fODF metrics in WM, AFDt scores similarly at 4%, AFD at about 9% and NuFO presents as the least reliable at 14%. Oddly, mean intra-subject variability is greater than inter-subject variability for this site for almost all quantities observed, except for fractional anisotropy (with the exception of the mean FA on DIAMOND fascicles), AFDt (in WM only) and free-water measurements, as well as for gamma distribution parameters excluding heiAD in WM and kappa in GM. With further inspections, it was noticed that for all measurements exhibiting this trend, two clusters of similar measurements could be formed, corresponding each to one of the sessions at which the subjects’ images were acquired, as presented in figure 15.

**Figure 15.**
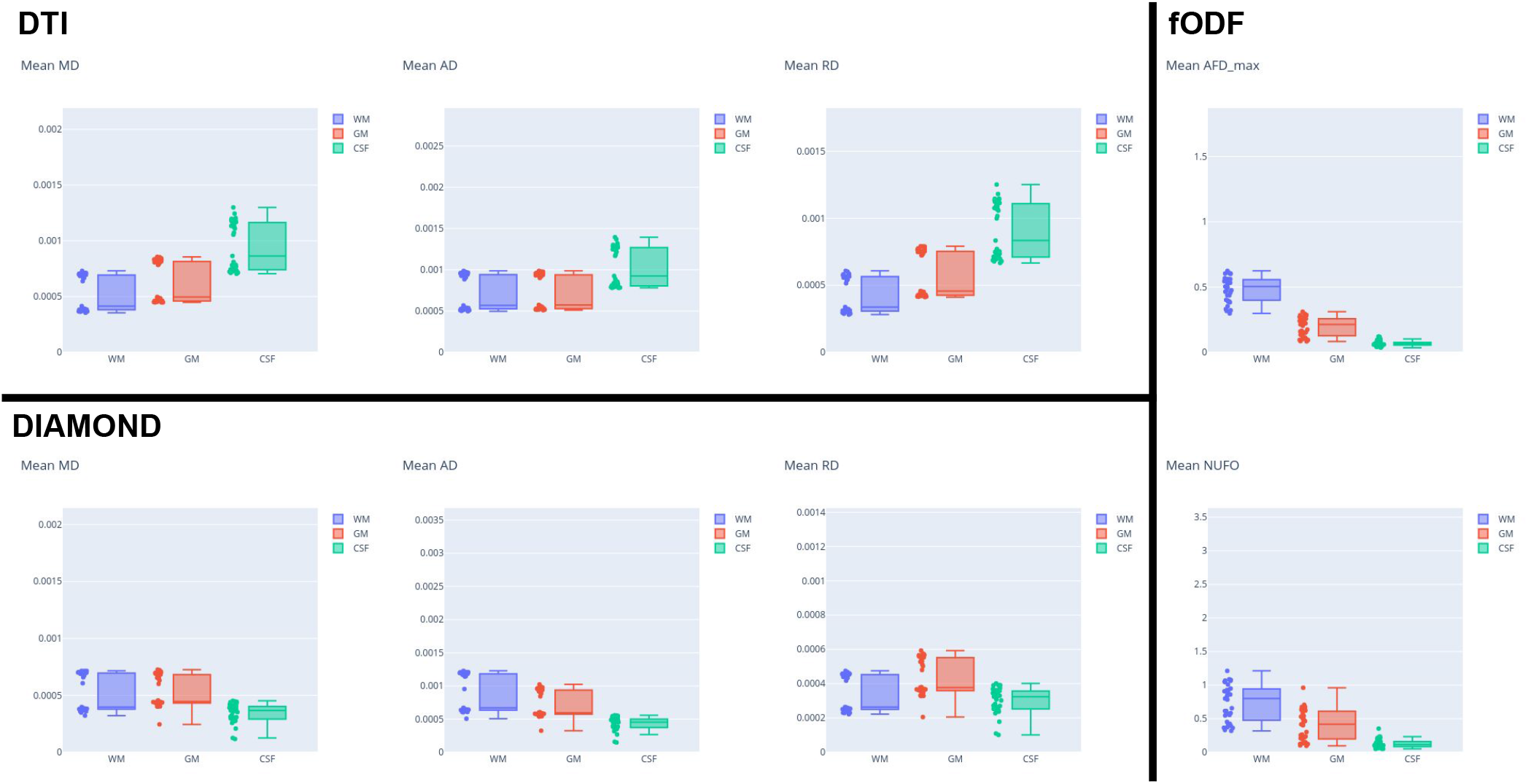
Clustering behaviour of the acquisition sessions of the subjects from the UC-Davis site.

Intra-vendor variability of Siemens scanners is computed on subjects from both Aix-Marseille and UC-Davis. For DTI metrics, variability in WM ranges from 12% to 14% for diffusivity measures and at around 8% for FA and is slightly lower in GM. For fODF, NuFO displays the highest variability (21% in WM, 32% in GM), followed by AFD (10% in WM, 22% in GM). Variability for AFDt is substantially lower at 4% in WM and 7% in GM. For measurements on the DIAMOND model in WM, MD is the least variable at 6% and fractional anisotropy scores at 10%, 2% bellow the fascicle based measure (max fFA). Results shows that performance from those measure is approximately two times more variable in GM. Free-water fraction is the most variable measure estimated at 28% in WM and 21% in GM. For gamma distribution parameters, values range widely depending on the parameter and on the tissue. For WM, kappa and kappaAD are the less variable at 7% and 10%, and hei and heiAD score at 33% and 18% respectively. For GM, the more variable parameter is kappaAD at 32%, followed by kappa and hei at around 17% and heiAD at 13%.

Inter-vendor variability is consistently higher than intra-vendor on all DTI measurements, with an increase in variability of 2% in FA and from 4% to 10% in diffusivity measures in both WM and GM. The same trend is observed for fODF metrics, AFDt displaying the less increase (2%) and NuFO the most (5%) in WM. For measures on the DIAMOND model, MD shows the greater increase (6% in WM and 3% in GM). FA and maxfFA variability remains stable, with a slight increase of 2% for the former and less than 1% for the latter. Free-water fraction variability increases substantially of about 5%. For gamma distribution parameters, a slight decrease of variability is observed for all parameters, except for hei which remains stable in WM and increases of about 3% in GM.

## 4 DISCUSSION

### 4.1 Reproducibility

Reproducibility is of prior importance when it comes to a good processing pipeline. It is a measure of how much a pipeline can, now and later in time, given the same data, configuration and underlying algorithm’s versions, output data that behaves the same and give similar results when analyzed. As for tractoflow [Theaud et al. (2020)], a novel pipeline for dMRI analysis of human subjects, this pipeline was designed with reproducibility in mind. Results displayed in table 2 confirm good reproducibility from DTI and fODF models computed, as well as from T1 registration processes (using ANTs registration) and tissues partial volume fractions computation (using ANTs Atropos). Scores are slightly lower for metrics estimated on the DIAMOND model, and significantly lower for gamma distribution parameters (kappa, kappaAD, hei, heiAD). This behaviour is expected since the DIAMOND implementation included in the pipeline does not allow to set the random number seed, which injects a quantity of variability in the outcomes of the algorithm. We would nonetheless advise against their usage in a statistical study as is, since they are bound to introduce a level of variability in the analysis. Note that this does not affect the measurements calculated (MD, FA, max fFA and fFW), since in the current implementation of the pipeline, their computation only rely on the mean tensor estimated on each fascicle, thus are independent of the gamma distribution parameters.

### 4.2 Pipeline implementation

In addition to providing a highly reproducible processing chain, our pipeline implementation also offers a high level of adaptability to the parametrization of its input’s acquisition sequences. Whilst designing the collection of processing modules, we took great care to make their configuration resilient to different spatial (voxel size, number of slices, orientation of slice and/or phase encoding) and orientational (number of b-values and/or gradient directions) configurations. This proved critical to adequately scale the parameter space of algorithms used by the pipeline, such as Topup, which rely on a definition of its deformation field given in millimetres in voxel space, and to allow disabling their execution when input requirements were unmet.

Nevertheless, we are aware the default values we offer could vary as algorithms are updated and that some use-case could benefit changing them. To that extent, is provided with most of them an additional configuration layer, presenting all possible parameters, to allow the end-user to finely adapt the pipeline to its specific study. We also allowed the user to disable all steps included in the pipeline, in the case one desires only to compute models and measures and has already preprocessed its data, or to speed up computation time by disabling steps deemed unnecessary after thorough quality control. Still, we do not recommend skipping any preprocessing steps we included in the pipeline when providing raw data from the scanner.

Furthermore, our implementation is ready for new use-cases for which our pipeline does not provide the adequate preprocessing and modeling steps. Using our modular design, a pipeline can easily be derived from ours, using new algorithms, workflows and/or input conventions, by adding the necessary objects in their respective module and binding their dataflow correctly.

### 4.3 Evaluation of variability

Lowering variability of measurements as much as possible is of prior importance in a study as it can destroy any predictive power of an analysis. In this study, we evaluated from the PRIME-DE database, using 3 sites that provided enough good quality DWI images, as well as T1w images, the variability of measurements provided by our pipeline on 4 levels : intra-subject for Sinai-Philips and UC-Davis, inter-subject for all 3 sites, intra-vendor for Siemens scanners (Aix-Marseille and UC-Davis) and inter-vendor, that is across all 3 sites.

Our results reflect common problems that still need to be addressed by the community to make MRI a fully reliable imaging modality. The increase of variability reported when including data from multiple scanner vendors and from multiple sites is representative of the need for data harmonization protocols capable of reducing the effects induced by the acquisition software and technology [Vollmar et al. (2010)][Tax et al. (2019)][Grech-Sollars et al. (2015)] as well as signal deviation occurring from environmental differences (temperature, humidity, time of day, etc.) [Book et al. (2021)][Meyer et al. (2016)]. Moreover, it cries at the need for consensus on acquisition protocols and to the creation and application of guidelines by the community, without which, group efforts like the PRIME-DE project cannot become fully relevant. As seen in table 1, acquisition parameters vary widely from one site to another, especially when it comes to the sampling of diffusion gradients’ directions. This has a direct impact on the validity of the reconstruction of diffusion models, since different b-values encodes different processes occurring in the imaged tissues [Veraart et al. (2019)][McKinnon et al. (2017)].

In our study, gradient sampling seems to have a limited impact on DTI measurements for Aix-Marseille, since shells with b-values higher than 1300 *mm*^2^*/s* are omitted. This however has a greater effect when comparing other sites to UC-Davis, since DTI was reconstructed for it from the only available shell at 1600 *mm*^2^*/s*, explaining the 3 to 4 fold increase in Siemens intra-vendor and global inter-vendor variability relative to the average inter-subject variability of associated sites. The impact becomes greater when looking at fODF and multi-tensor reconstruction (DIAMOND), since those model make use of all b-values. This makes Aix-Marseille inter-subject variability increase since the analysis becomes a comparison between 2 subjects acquired with single-shell sampling, and 2 others with multi-shell, the latter presenting a richer portrait of the diffusion process by the usage of higher gradient weighting. Both fODF and multi-tensor reconstruction are known to benefit from the higher b-values [Yang et al. (2015)][Taquet et al. (2015)][Scherrer and Warfield (2012)][Scherrer and Warfield (2010)]; the quantities estimated from those models are destined to vary from the case of a low b-value single-shell acquisition.

Another trend observed in our study hinting to the need of guidelines and protocols is the high mean intra-subject variability observed for UC-Davis. While analyzing this site, we noticed a strong discrepancy of diffusion measurements between the two sessions images were acquired at. This behaviour could be the result of equipment upgrades, software or hardware, as well as poor sequence configuration, resulting in an ill-conditioning of the tissues required for diffusion acquisition. The only quantities were this behaviour was not detected are fractional anisotropies (FA and max fFA) and total apparent fiber density (AFDt). Nevertheless, this makes the usage of separate sessions from UC-Davis impossible in a study of diffusion measurements as the variability would erase any statistical power of an analysis. We hope in the future, good quality control could be integrated in the scanners to prevent or correct the acquisition of images presenting this kind of behaviour.

## 5 LIMITATIONS AND FUTURE WORK

In the current version of our pipeline, we implemented a robust and efficient preprocessing chain of state-of-the-art algorithms to correct for well know artifacts affecting the MRI signal, such as background noise, gibbs ringing, susceptibility distortions, motion, signal dropout and intensity disparities within tissues. We are well aware that our technique is not universal and that some use-cases may require the execution of different algorithms than the one we considered. MRI preprocessing still need to be thoroughly validated; the effect of applying specific techniques and algorithms, as well as their ordering relative to one another, requires to be quantified in order to attain an adequate level of consensus in the community. We designed our pipeline and its modular framework taking those facts into account, to ease the process of swapping and upgrading its different steps, in order to fit specific study cases. We also plan to release in the near future a version of the pipeline integrating alternative denoising techniques that are popular in the diffusion MRI research community based on recommendations from the ISMRM diffusion study consensus effort, once their work becomes available. Inasmuch, we intent to include brain extraction workflows, for both anatomical and diffusion weighted images, adapted to the morphology of input subjects. Good solutions have been developed that display good resiliency on human subjects, but their extension to other primates and to small animals is still to be validated and can lead to segmentation lacking specificity. As the field continues to evolve and those issues become addressed, we will consider the addition of those solutions in our pipeline.

Moreover, for our implementation, we chose a limited subset of 3 local models to represent the diffusion process in dMRI voxels. More algorithms, especially in the category of multi-tensor models, could be considered in the future, such as the ball and stick [Behrens et al. (2003)][Behrens et al. (2007)][Jbabdi et al. (2012)], ball and zeppelin [Sotiropoulos et al. (2016)] models and NODDI [Zhang et al. (2012)]. The inclusion of Diffusion Kurtosis Imaging [Jensen et al. (2005)] could also be considered. More general models of the Ensemble Averaged Propagator (EAP) could also be added, using for example the Simple Harmonic Oscillator Based Reconstruction and Estimation (3D-SHORE) or the Mean Apparent Propagator (MAP) [Ö zarslan et al. (2013)], along with relevant measurements computed on them. Furthermore, we plan to release supplemental modules to compute tractograms upon the computed models the pipeline offers.

It is part of future work to adress better computing resources management, there is room for improvement. The Nextflow scripting language offers extensive capabilities to define per process resources prerequisites. Using them, we were able to define clearly the number of processing cores (CPU) and need for hardware accelerators (GPU) for all processes in our library and ensure a high level of parallelization. However, we do not yet control for requirements in memory (RAM) of each individual process executed. The lack of specification for memory usage of multiple algorithms we used, combined with the difficulty of obtaining the actual memory space of compressed NIFTI images using the Nextflow language makes it hard to define a clear requirement to respect at execution time. Specifying an overestimated precondition on memory needs could lead to a fewer number of processes executed at the same time, which would impede the parallelization capabilities of the pipeline and result in a waste of computing resources. We are aware that this behaviour could lead to execution problems, more so when processing high resolution images that spans multiple gigabytes on disk. To mitigate the effects of that limitation, we opted for a retry strategy for processes failing for that reason, combined with a restriction of the number of parallel executions of memory hungry processes.

## 6 CONCLUSION

In this work, we built a reliable and reproducible pipeline, able to tackle the task of preprocessing and computing diffusion models on multi-resolution diffusion MRI data. We also presented our modular library of Nextflow processes and workflows using state-of-the-art and cutting edge DWI processing technologies, designed to be easily upgradable and adaptable. We used our pipeline to analyze the variability of 3 sites (32 subjects) included in the PRIME-DE database presenting good quality DWI data. We showed that even if promising, that data exhibited a great level of variability and its usage should be done with care to prevent instilling uncertainty in statistical analyses. This is a good example of the effects of the lack of consensus in the diffusion MRI field when it comes to specifying guidelines for acquisition procedures.

The hurdle of defining a gold standard in dMRI makes this problem even more difficult to overpass, but a level of standardization and harmonization must be attained in order to reduce variability in imaging between subjects, scanner and sites as much as possible. Without it, conjugate efforts like the PRIME-DE become less relevant for the study of the brain and its properties. It is through conjoint endeavours such as the consensus effort from the ISMRM diffusion study group and others, and by the development of reproducible, robust and maintainable software, that we as a community will make of diffusion MRI a reliable modality.

## Supporting information

Default configuration file for mrHARDIflow

## DATA AVAILABILITY STATEMENT

The pipeline developed for this study and the nextflow modules library are available online at github.com/boiteaclou/mrHARDIflow. The processed diffusion data, including computed models and diffusion metrics, are available on Zenodo at doi.org/10.5281/zenodo.5719453.

