## Supplementary material for "mrHARDIflow : A pipeline tailored for the preprocessing and analysis of Multi-Resolution High Angular diffusion MRI and its application to a variability study of the PRIME-DE database": Default configuration file for mrHARDIflow

// Inputs declaration
params.default_readout = false                  // Default readout time of input images (used when none supplied with the DWI image)
params.default_multiband_factor = false         // If supplied, specifies as default that slices of input images were acquired with that multiband factor (used when none supplied with the DWI image)
params.default_is_interleaved = null            // If true, specifies as default that slices of input images were acquired interleaved (used when none supplied with the DWI image)
params.default_slicing_direction = false        // Default slice stacking direction of input images (used when none supplied with the DWI image)
params.default_phase_direction = false          // Default phase encoding direction of input images (used when none supplied with the DWI image)

// Use gpu if available
params.use_cuda = true                          // If true, CUDA will be used (Requires one or more Nvidia GPU)

// Resources conservation
params.max_cpu_per_process = null               // If supplied, will limit the maximal number of cpu cores a task can use
params.max_attempts = 3                         // Maximal number of attemps of a processing task before failing


// Enable and disable pipeline steps

// Global processes
params.resample_data = true                     // If true, adds a final resampling step to the T1, b0, DWI, masks and segmentations at the end of preprocessing 

// T1 processes
params.denoise_t1 = true                        // If true, performs background noise correction on the T1 image
params.t1_intensity_normalization = true        // If true, performs N4 normalization on the T1 image (Don't use if the data has been reshuffled)
params.t1mask2dwi_registration = true           // If true, registers the T1 mask the the DWI space instead of using a BET mask
params.register_t12b0_denoised = true           // If true, in adition to the registration after Topup, an aditional registration step will be insterted at the end of the preprocessing
params.register_syn_t12b0 = true                // If true, adds a SyN deformation field to the T1 to b0 registration

// Segmentation processes
params.generate_tissue_segmentation = true      // If true, generates a tissue segmentation (WM, GM, CSF) from the T1 image
params.generate_wm_segmentation = true          // If true, generates a white matter segmentation from the T1 image

// DWI processes
params.gaussian_noise_correction = true         // If true, performs background noise correction (MP-PCA) on the DWI
params.gibbs_ringing_correction = false         // If true, performs gibbs rigging correction (Don't use if the data was acquired using partial fourier)
params.normalize_inter_b0 = true                // If true, performs normalization between b0 volumes of both phase acquired DWI
params.topup_correction = true                  // If true, performs suceptibility correction on the DWI
params.eddy_correction = true                   // If true, performs eddy currents and motion correction on the DWI
params.dwi_intensity_normalization = true       // If true, performs N4 normalization on the DWI

// Reconstruction processeses
params.recons_dti = true                        // If true, reconstructs the single tensor model (DTI)
params.recons_csd = true                        // If true, reconstructs the fODF model using CSD (SSST or MSMT)
params.recons_diamond = true                    // If true, reconstructs the DIAMOND multi-tensor model

// WORKFLOWS PARAMETRIZATION

// Global preprocessing parameters
params.resampling_resolution = false            // If a float number is supplied, T1, b0, DWI, masks and segmentations will be resampled to this isotropic voxel size after preprocessing
params.force_resampling_sequential = true       // Resampling can be a memory hungry operation. If true, forces the execution of resampling processes to be sequential
params.b0_threshold = false                     // If supplied, will use this value as a upper threshold to determine b0 volumes

// T1 preprocessing parameters
params.register_syn_t12b0_with_mask = true      // Use brain masks when computing SyN deformation of T1 to b0

params.nlmeans_t1 = true                        // If true, performs Non-Local Means Rician bias denoising on the T1 image. Else, uses Gaussian denoising

// DWI preprocessing parameters

// B0 normalization
params.b0_normalization_strategy = "linear"     // Normalization strategy when adjusting the mean of b0 volumes in forward and reverse acquisition. Three modes are
                                                // available to describe how to behave with blocks of diffusion volumes between 2 blocks of b0 :
                                                // - linear : perform linear interpolation of the mean value of b0 before and after the block of diffusion with weighting
                                                //            based on the distance the a b0 in the sequence
                                                // - first : use the block of b0 coming before a block of diffusion
                                                // - last : use the block of b0 coming after a block of diffusion

// Eddy
params.eddy_select_gpu = true                   // Perform GPU selection when running Eddy (required if multiple GPU are available on a single computing node)
params.cuda_max_parallel = 4                    // Maximal number of parallel execution of CUDA processes

// Bet
params.bet_f = 0.5                              // f value for BET computation

// DTI
params.max_dti_bvalue = 1300                    // Maximal b-value threshold for DTI reconstruction

// CSD
params.msmt_odf = false                         // Perform Multi-Shell Multi-Tissue fODF reconstruction if possible
params.frf_fa = 0.75                            // Initial search value for single-fiber population voxels
params.frf_min_fa = 0.55                        // Minimal FA of single-fiber population voxels
params.frf_min_nvox = 200                       // Minimal number of voxels to select to perform fiber response estimation
params.frf_radii = false                        // Research cube radii for fiber response estimation
params.frf_center = false                       // If specified (voxel index as a list of 3 integers), will search for voxels for fiber response estimation from this point
params.max_fa_ventricle = 0.15                  // Maximal FA value in ventricles for fODF peak threasholding
params.min_md_ventricle = 0.0012                // Minimal MD value in ventricles for fODF peak threasholding
params.ventricles_center = false                // If specified (voxel index as a list of 3 integers), will perform ventricles detection from this point
params.fodf_max_absolute_factor = 2.0           // Absolute threshold for fODF peak extraction
params.fodf_relative_thr = 0.2                  // Relative threshold for fODF peak extraction
params.sh_order = 8                             // Use spherical harmonics order 8 to reconstruct fODF

// DIAMOND
params.n_fascicles = 3                          // Maximal number of fascicles (tensors related to fiber bundles) to estimate in each voxel
params.fascicle_model = "diamondNCcyl"          // Model for the fascicle representation and gamma distribution parametrization
params.model_selection_with_tensor = false      // Do model selection using full tensor instead of the principal direction only
params.estimate_restriction = false             // Estimate restriction (extra-axonal fraction and diffusivity) for each fascicle
params.restriction_tensor = false               // Use a tensor to represent extra-axonal diffusivity
params.normalized_fractions = true              // Normalize volume fraction for fascicles and free water in voxels to 1
params.free_water_tensor = false                // Use tensor to estimate free water diffusivity

// Miscelaneous
params.random_seed = 1234                       // Random seed for stochastic algorithms
